# The Cannibalistic Trade-Off: Why Human Cannibalism Emerges and Why Taboos Suppress It

**DOI:** 10.64898/2026.02.10.705014

**Authors:** Michal Misiak, Petr Tureček

## Abstract

Cannibalism is among the strongest and most widespread food taboos in human societies, yet archaeological, ethnographic, and historical evidence indicates that it has repeatedly emerged across diverse human populations. This coexistence of recurrent practice and persistent prohibition raises a fundamental question: when does cannibalism become adaptive, and why is it typically suppressed? We address this problem using a formal model that treats cannibalism as a potential food source subject to energetic benefits and multiple sources of cost.

Nutritional gains are modelled using a saturating function of caloric intake, while costs arise from acquisition, digestion, and infection. Infection costs are represented as a stochastic process whose mean increases with the length of the trophic transmission chain, capturing the risks associated with repeated within-species consumption. Analysing the expected energetic balance across levels of food availability and cannibalism order reveals narrow ecological conditions in which cannibalism yields a positive expected balance and broader conditions in which it is strongly disfavoured.

The model provides a framework for interpreting archaeological and ethnographic findings by specifying boundary conditions and identifying the most probable ecological scenarios under which different forms of cannibalism are expected to occur. The results predict that cannibalism is most likely under extreme resource scarcity, when acquisition costs are low and infection risks are constrained, while sustained or high-order cannibalism rapidly becomes unviable due to escalating infection costs. Overall, the findings suggest that cannibalism is best understood as a conditional trade-off rather than a behavioural anomaly, with cultural taboos functioning as adaptive responses to nonlinear epidemiological risks.

## 1. Introduction

Cannibalism is subject to one of the most pervasive taboos (Macbeth et al., 2007). Historically, this cultural prohibition is so pronounced that only a small number of societies have been reported to engage in highly restricted ritual forms of the practice, and even these accounts are often contested. Despite this strong and morally charged taboo (McHugh et al., 2017), archaeological, ethnographic, and genetic evidence indicates that cannibalism has nevertheless emerged repeatedly across diverse human populations (Bello, 2024; Byard, 2023; Mead et al., 2003). If cannibalism occurred often enough to leave archaeological and genetic traces, why is it so consistently constrained or suppressed across cultures? These observations suggest that cannibalism may involve trade-offs that allow it to occur under some conditions while limiting its persistence.

### 1.1. Human Cannibalism

Cannibalism refers to the consumption of human flesh by other humans. Evidence for cannibalistic behaviour appears repeatedly across human history, sometimes emerging under extreme conditions of starvation (Byard, 2023; Petrinovich, 2000; Schutt, 2018). It has also been documented in isolated clinical cases involving individuals, often discussed in psychiatric or forensic contexts (Oldak et al., 2023). In this paper, we do not primarily address such isolated incidents, which typically involve the violation of strong cultural taboos and entail heavy social sanctions within the societies in which they occur. Instead, we focus on cannibalism as a population-level subsistence practice that is socially permitted.

Evidence for human cannibalism spans a broad temporal, geographic, and cultural range (Lindenbaum, 2004). Archaeological assemblages document anthropogenic modification of human remains consistent with butchery, marrow extraction, and cooking in multiple prehistoric contexts, indicating that cannibalism has occurred repeatedly throughout human evolution rather than being confined to a single period or region (Bello, 2024; Saladié et al., 2015). Some researchers have argued that in certain prehistoric populations, such as Magdalenian groups, cannibalism may have been widespread and relatively common, forming a recurrent component of social or subsistence practices (Marginedas et al., 2025). There are also interpretations proposing similarly recurrent cannibalistic practices in earlier *Homo* populations (Carbonell et al., 2010). At the same time, evidence from Neanderthal assemblages has often been interpreted as indicating more episodic cannibalism, frequently associated with periods of resource scarcity, highlighting substantial variation across hominin populations (Yustos et al., 2015).

Anthropological research commonly distinguishes between endocannibalism and exocannibalism, two forms that differ in social context and mode of acquisition (Bello, 2024; Lindenbaum, 2004). Endocannibalism refers to the consumption of members of one’s own social group, most often in mortuary contexts. Well-known example include the Fore of New Guinea, who became the focus of medical investigation following the emergence of a rare prion disease, kuru, associated with their mortuary practices (Liberski et al., 2012; Mathews et al., 1968). Exocannibalism, by contrast, involves the consumption of individuals from outside the social group, typically enemies killed during raids, ambushes, or warfare. For example, Māori warfare cannibalism has been interpreted as a practice aimed at humiliating defeated opponents (Moon, 2008). While both forms involve the consumption of human flesh, they differ substantially in their acquisition costs, as exocannibalism entails the risks associated with hunting and killing another human (Byard, 2023). Importantly, these forms are not mutually exclusive: ethnographic evidence indicates that some populations practiced both endocannibalism and exocannibalism, as documented among the Wari’ (Vilaça, 2005).

Ethnographic evidence for cannibalism is frequently contested. Many reports were based on second-hand accounts and have been argued to reflect colonial narratives that exaggerated accusations of cannibalism to stigmatise Indigenous populations and justify their subjugation (Arens, 1979; Pickering, 2017). Historical sources, however, document highly regulated forms of European medicinal cannibalism, which operated within strict cultural rules governing who could be consumed, in what form, and for what purposes (Sugg, 2015). For example, in the seventeenth century in parts of the Germanic world, there was a practice of drinking the blood of executed men as a treatment for epilepsy (Sugg, 2015). Today, the cultural prohibition against cannibalism is so strong that only a small number of groups are reported to engage in its highly restricted ritual forms, and even these accounts are interpreted as performances for tourists (e.g., the Aghori in India or the Korowai in Papua).

### 1.2. Benefits and Costs Associated with Cannibalism

From the perspective of human behavioural ecology, cannibalism can be examined as a form of food acquisition within frameworks such as optimal foraging theory (Rodríguez et al., 2019). This approach builds on principles from animal behavioural ecology and evolutionary theory, treating human behaviour as shaped by natural selection and constrained by factors that influence fitness, including energy availability (Winterhalder & Smith, 2000). Because energetic intake is a key limiting factor for survival and reproduction, a central assumption of this approach is that foragers tend to favour behaviours that maximise nutritional returns relative to associated costs. From this perspective, cannibalism can be analysed alongside other subsistence choices, with human bodies treated as potential resources subject to the same cost–benefit considerations as other food items (Rodríguez et al., 2019).

Archaeological evidence suggests that human bodies were sometimes processed and consumed in ways consistent with other prey, following similar cost–benefit logic (Rodríguez et al., 2019). However, humans rank relatively low among potential prey in terms of energetic returns, and human flesh offers no exceptional caloric advantage compared to similarly sized animals, and is nutritionally inferior to large game (Cole, 2017). These findings imply that cannibalism is unlikely to be explained by energetic benefits alone and would be favoured only under restricted ecological conditions, such as when higher-yield resources are scarce. While subsistence-based approaches help clarify when cannibalism may be nutritionally plausible, they do not account for additional costs that can accumulate across repeated consumption events.

The costs may substantially limit viability of cannibalism as a subsistence strategy. These include typical food acquisition costs linked to digestion, locating, killing, or processing meat (Kraft et al., 2021).

A central cost of cannibalism, however, arises from its role as a route of disease transmission (Rudolf & Antonovics, 2007). When humans consume herbivores such as deer or cattle, pathogens carried by these animals typically face substantial barriers to establishing infection in the human body. The risk increases when humans consume carnivores, because predators accumulate pathogens through repeated consumption of prey over their lifetime. There are many pathogens and parasites that specialize in transmission via the trophic route along the food chain (Lafferty, 1999; Park, 2019). Some of them (such as *Trichinella spiralis*) do not map their life-cycle stages onto fixed species and can be passaged indefinitely between many different predators and scavengers (Pozio et al., 2009). Cannibalism represents the extreme opportunity for such pathogens: when humans consume other humans, pathogens are transmitted between individuals with identical physiology, maximising the likelihood of successful infection and subsequent spread. For humans, this risk is even greater than for most other predators, because cannibalistic practices often involve multiple individuals consuming the same conspecific, allowing a single infected body to expose several members of the group simultaneously and thereby increasing the risk of community-wide transmission (Rudolf & Antonovics, 2007).

A particularly clear empirical illustration of the epidemiological costs associated with cannibalism is provided by the kuru epidemic among the Fore of Papua New Guinea (Klitzman et al., 1985). Kuru emerged as a result of endocannibalistic mortuary practices, in which deceased relatives were consumed as part of culturally sanctioned funeral rituals (Liberski et al., 2012). The disease was invariably fatal and transmitted through prions, infectious protein particles that are exceptionally resistant to heat and therefore not eliminated by cooking. As a result, standard food-processing practices that reduce infection risk for many pathogens were ineffective in this case. The epidemic persisted for decades and declined only after the abandonment of cannibalism, despite no corresponding changes in caloric availability or broader subsistence strategies (Alpers, 2008).

Evidence from human evolutionary genetics suggests that the Fore case is not an isolated anomaly, but part of a broader pattern of recurrent epidemiological costs associated with cannibalism. Genetic evidence complements archaeological findings by showing that cannibalism has carried substantial fitness costs over human evolutionary history. In particular, signatures of long-term balancing selection at the human prion protein gene suggest repeated exposure to transmissible neurodegenerative diseases in prehistoric populations, plausibly linked to cannibalistic practices (Mead et al., 2003). This indicates that cannibalism occurred often enough, and with serious enough consequences, to leave a detectable genetic trace.

### 1.3. The Cannibalistic Trade-Off

Cannibalism has emerged repeatedly across human history, yet it has been strongly tabooed in both historical and contemporary societies (Byard, 2023; Petrinovich, 2000; Schutt, 2018). Although cannibalism can provide nutritional benefits, it also entails costs, including acquisition risks, digestion, and most critically, infection risks that may escalate with repeated consumption.

Existing approaches address these components in isolation. Subsistence-oriented models emphasise energetic returns (Cole, 2017; Rodríguez et al., 2019), whereas epidemiological work highlights the pathogen risks (Rudolf & Antonovics, 2007). Here, we conceptualise cannibalism as a trade-off between nutritional gains and accumulating costs. To examine this trade-off formally, we develop a model that integrates nutritional benefits with acquisition, digestion, and infection costs, allowing us to identify the ecological and epidemiological conditions under which cannibalism becomes viable and those under which it is disfavoured.

Human cannibalism cannot be fully understood by analogy to cannibalism in other animals (Claessen et al., 2004; Fox, 1975; Rudolf, 2007). Human subsistence behaviours are highly flexible and shaped by culturally transmitted rules, technologies, and taboos. This includes distinct forms of cannibalism, such as endocannibalism and exocannibalism, as well as the use of fire and food-processing practices that can reduce infection risk (Smith et al., 2015). Together, these cultural and technological factors fundamentally change the cost–benefit structure of cannibalism, making a specifically human-focused framework necessary.

## 2. Methods

We evaluated the expected energetic balance associated with incorporating different food sources across a two-dimensional parameter space defined by background caloric intake and cannibalism chain length. For each parameter combination, we identified threshold conditions corresponding to zero net energetic balance and visualised these thresholds as contours on the parameter plane.

Analyses were conducted under four scenarios defined by the factorial combination of two acquisition cost regimes (hunting versus food preparation only) and two pathogen load conditions (raw versus heat-processed meat). All computations and visualisations were performed in R (version 4.0.3). The full set of functions used in the analyses is available online (https://osf.io/eyvs3).

### 2.1. Formal Model

We have constructed a mathematical model guiding the inclusion of a new nutrition source into a diet. Food source profitability is a function of its Caloric value (*n*), the total energy available from other previously incorporated food sources (*x*), the maximum amount of effective Calories (*ϕ*), number of Calories with 100% effectiveness (*ψ*), and expected costs linked to the potential new source of nutrition (*c*).

We parcelled the costs into three independent components: digestion costs (*c*_*D*_, characterised by a low median and low variance), acquisition costs (*c*_*A*_, with very diverse medians and variances depending on the food source), and infection costs (*c*_*I*_, with a low median - unless carrion is the considered food source - and variance proportional to the longest undisturbed trophic chain leading to the source.) We treat the total cost as a sum of expected values of distributions attributed to said components (*c* = 𝔼[*c*_*D*_] + 𝔼[*c*_*A*_] + 𝔼[*c*_*I*_]). This mathematical model provides a theoretical foundation to judge many intuitions and claims surrounding human subsistence strategies, including occasional, facultative, and customary cannibalism.

#### 2.1.1. Mathematical Model of Food Source Utilisation

The saturation function governs energetic benefits in the model. The saturation and benefit functions are illustrated in Figure 1 using a hypothetical food portion. We employ the concave part of a sigmoid curve, specifically a logistic function

**Figure 1.**
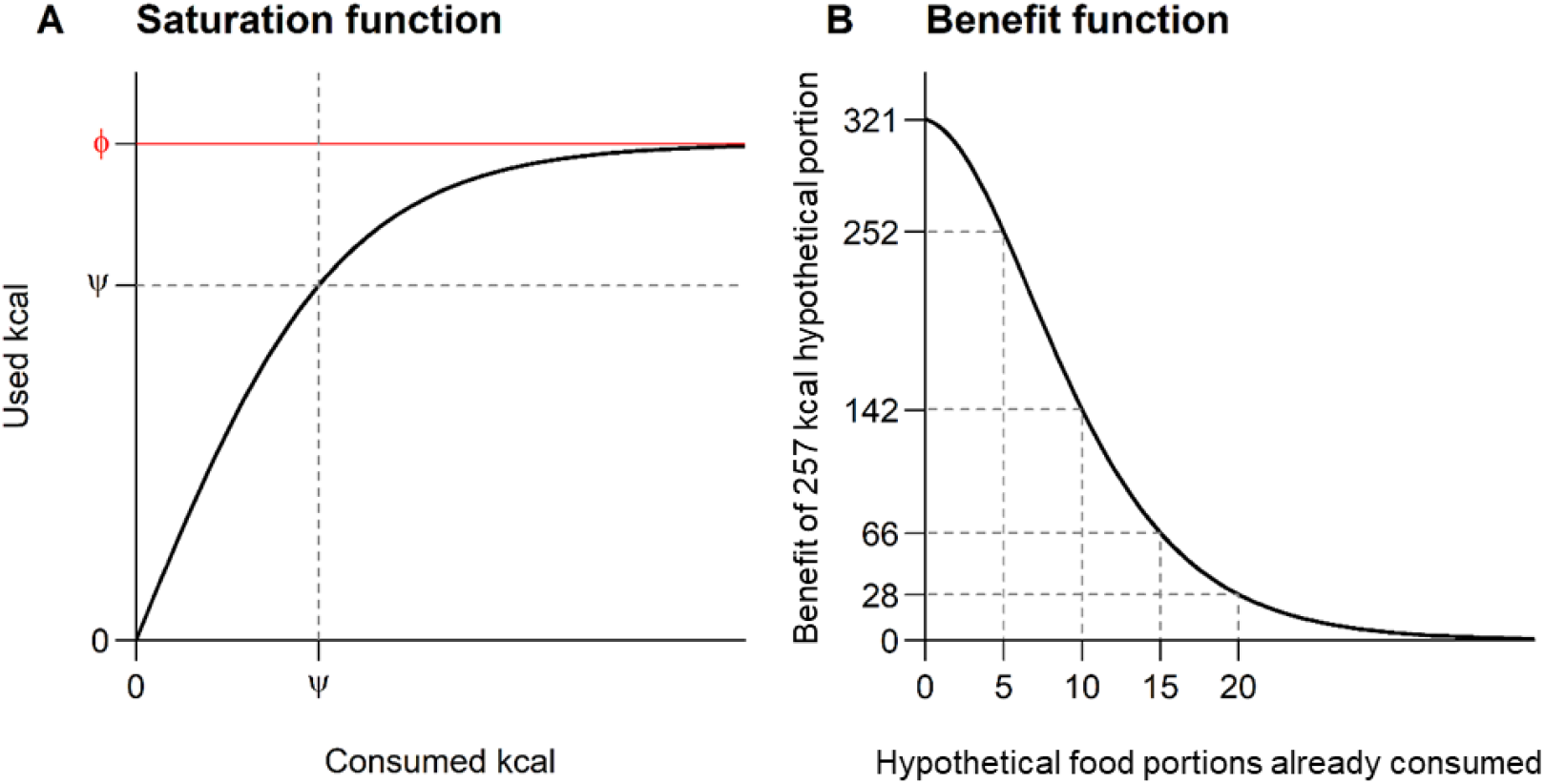
Generic saturation function scaled by parameters ϕ and ψ (A) and the corresponding benefit function describing the effective caloric contribution of a hypothetical food portion after x calories have already been consumed (B) *Note. ψ* = 2500 (recommended daily Caloric intake of an adult human male), *ϕ* = 3500 (his basal metabolic rate multiplied by 1.9), and *n* = 287*kcal* (illustrative food portion).

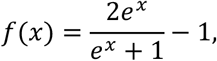

which is subsequently scaled using parameters *ϕ* and *ψ* as

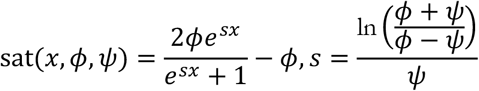

This parametrisation ensures that the saturation function approaches an upper bound of *ϕ* and satisfies the condition sat(*ψ*) = *ψ* (Figure 1A; see Supplement S1 for details).

The expected caloric benefit of a food source of size *n* calories, following prior incorporation of food sources with a combined caloric value *x*, is defined as

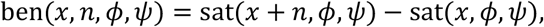

After algebraic simplification (Figure 1B; Supplement S2), this expression can be written as

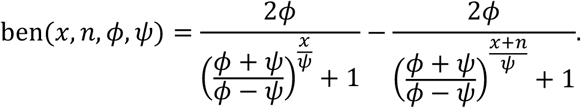

Each cost component is characterised by a log-normal distribution. This distribution satisfies a maximum-entropy criterion for strictly positive random variables specified by two parameters, such as the mean and variance (alternatively, the mean and the mean logarithm). Log-normal distributions naturally arise from random multiplicative processes, in which values are generated as the product of a central tendency and stochastic multiplicative factors; under this formulation, a factor *e*^*x*^ is as likely as its reciprocal *e*^−*x*^.

A log-normal distribution is conventionally defined as *e*^𝒩(*μ,σ*)^, where 𝒩(*μ, σ*) denotes a normal distribution with mean *μ* and standard deviation *σ*. Parameter *μ* corresponds to the natural logarithm of the distribution median (*M* = *e*^*μ*^), while *σ* quantifies the magnitude of multiplicative stochastic variation acting on this median. The expected value of a random variable *X* drawn from a log-normal distribution *e*^𝒩(*μ,σ*)^ is given by 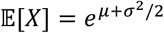. To avoid ambiguity, we denote the arithmetic mean (equal to the expected value) by *μ*_*X*_ and the corresponding standard deviation by *σ*_*X*_.

Unlike the normal distribution, the expected value of a log-normal distribution depends jointly on both the median and the degree of stochasticity, while the median itself remains unchanged. This property reflects the asymmetry of multiplicative processes acting on strictly positive quantities: random variation cannot reduce values below zero but can generate arbitrarily large outcomes.

The total expected balance associated with incorporating a potential new food source is defined as the difference between expected energetic benefits and expected costs. Accordingly, the balance function is given by

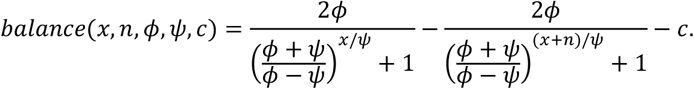

In principle, the cost term in the balance function may incorporate additional costs or benefits (represented as negative costs) that are independent of the saturation function, such as benefits associated with costly signalling, conspicuous consumption, or reciprocity. In the present analysis, however, we restrict attention to three primary cost components: digestion costs, acquisition costs, and infection costs.

Digestion costs are unavoidable and apply to all food sources, regardless of how low acquisition or infection costs may be. These costs reflect energetic expenditures associated with physiological processing of food and impose an upper bound on consumption, even under conditions of abundance.

Acquisition costs capture the energetic expenditures required to obtain and prepare a food source. More specifically, the ratio between expected acquisition costs 𝔼[*c*_*A*_] and expected nutritional value *n* influences whether a given food source is incorporated into the active diet. Acquisition costs include the energy required to locate, capture, and process units of the food source.

Infection costs reflect energetic losses associated with morbidity or mortality risk following consumption. In the absence of such costs, consumption of conspecific carcasses would be expected to be common; however, this behaviour is rarely observed in humans or other animals. Infection costs also account for the tendency of most carnivores to preferentially consume freshly killed, apparently healthy prey rather than carcasses or visibly diseased individuals.

The parameters of the infection cost distribution can be expressed in terms of pathogen load and trophic chain length. Let infection costs attributable to a single source of pathogenicity be distributed as 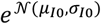. The expected value of this distribution is a small positive real number, so both *μ*_*I*0_ and *σ*_*I*0_ are bound to be small. When multiple independent pathogenic sources are present, their cumulative effect is assumed to follow a multiplicative rather than an additive process, because they may amplify or counteract each other. The overall pathogenicity of the prey is, however, expected to grow with the number of unit pathogens. Therefore, if 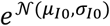 describes infection costs associated with a single pathogen, the cumulative effect of two pathogens is given by the product of two such distributions, which is equivalent to 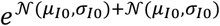.

For two uncorrelated random variables *X*_1_ and *X*_2_ with means 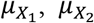 and standard deviations 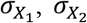, the distribution of their sum has mean 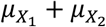 and standard deviation 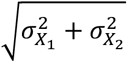, as variances – rather than standard deviations – are additive (Fisher, 1918).

Infection costs are therefore characterised by a log-normal distribution 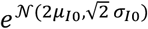for two unit pathogens, and by 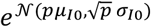for *p* unit pathogens. Under the same scaling logic, increasing pathogen load by a factor of one hundred corresponds to a hundredfold increase in the location parameter *μ*, but only a tenfold increase in the dispersion parameter *σ*, yielding a distribution 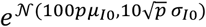. This property allows the distribution to be rescaled according to relative pathogen load without requiring explicit specification of *μ*_*I*0_or *σ*_*I*0_. More generally, changing pathogen load by a factor *ζ* transforms the infection cost distribution 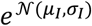 to 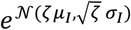.

Variation in overall pathogenicity further permits incorporation of phylogenetic relatedness between predator and prey into the model. Greater phylogenetic proximity implies a higher proportion of *p* prey-associated pathogens that remain viable in the consumer. Consequently, when humans are the consumers, infection costs are expected to be lower for raw fish than for raw mammals, and highest for consumption of conspecifics (see Supplement S7 for illustrative parameterisations).

Trophic chain length does not alter the median of the infection cost distribution, as the total pathogen load is assumed to remain constant. However, it affects the variance of infection costs. Each of the *p* prey-associated pathogens that remain viable in the consumer may undergo random changes in pathogenicity during trophic transmission, with some becoming less harmful and others more harmful. We assume that variation in pathogen load or strain composition across tokens of the same food source is negligible, and that all variability in infection costs per pathogen (*σ*_*I*0_) arises from stochastic changes in pathogenicity during transmission. This assumption allows changes in expected infection costs to be approximated as a function of trophic chain length without introducing additional parameters to distinguish mutation rates from baseline variation.

Accordingly, if 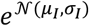 characterises infection costs for a trophic chain length of *ρ* = 1, then infection costs for *ρ* = 2 are given by 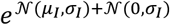. For an arbitrary trophic chain length *ρ*, infection costs can be expressed as

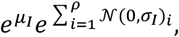

which is equivalent to 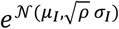.

Throughout the remainder of the article, we refer to *ρ* as the order of cannibalism, thereby abstracting away from cases in which lower-order cannibals consume animal carnivores. Although the widespread avoidance of top predator meat may be partly attributable to elevated pathogen loads, the present analysis focuses on the consumption of conspecifics. Accordingly, phylogenetic relatedness to the prey—captured by the baseline parameters *μ*_*I*_ and *σ*_*I*_ associated with first-order cannibalism—is treated as more consequential than the prey’s position in the non-cannibalistic trophic chain.

Parameter *ρ* becomes particularly important in trophic chains composed exclusively of members of a single species. In such cases, trophic chain length is not bounded a priori, and pathogens adapted to both consumer and consumee remain highly viable across successive transmission events. Even in the absence of an explicit model component capturing selection for pathogens that spread efficiently through cannibalistic trophic chains, expected infection costs increase exponentially with the order of cannibalism, such that

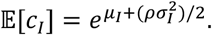

An illustrative simulation of 50 stochastic trophic chains demonstrates the relationship between the number of pathogenicity-altering events and the resulting shape of the log-normal infection cost distribution (Figure 2). Consistent with analytical expectations, the median of the distribution remains unchanged across trophic chain lengths, whereas the arithmetic mean, corresponding to the expected value, increases as a consequence of rare but influential chains in which a small number of cumulative changes generate disproportionately high pathogen virulence.

**Figure 2.**
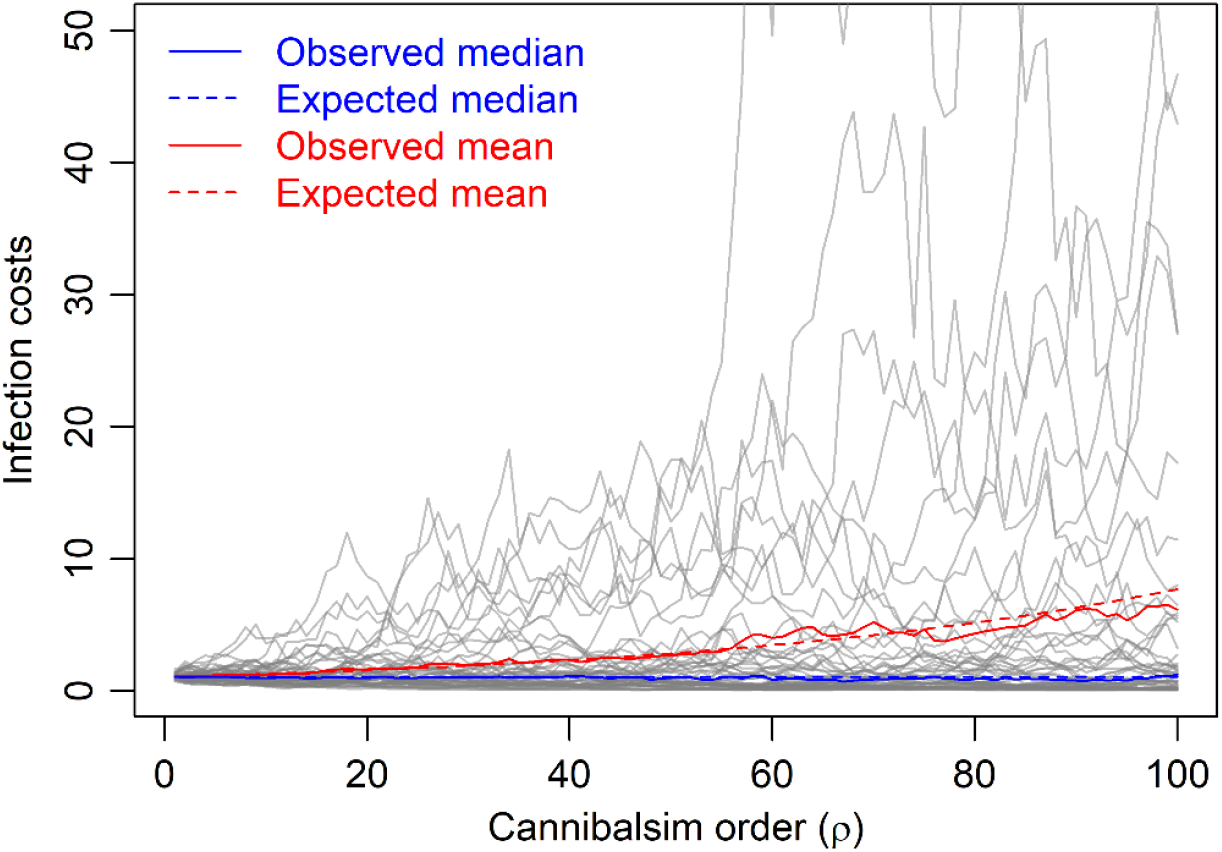
Results of a computer simulation showing the changes in the expected value with the number of random changes in pathogenicity. *Note*. Fifty trophic chains were simulated, each grey line corresponds to a single trophic chain, *μ*_*I*_ = 0.04, *σ*_*I*_ = 0.2

As outlined above, human flesh is not expected to constitute a favourable food source under conditions of abundant alternative resources. However, when nutritionally adequate and readily accessible food sources become scarce, the model predicts that consumption of conspecifics may yield a positive energetic balance, particularly when the order of cannibalism *ρ* is constrained. Using the balance function, we solve for the caloric threshold *t*_0_ defined by

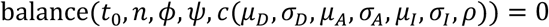

which identifies the level of background caloric intake at which a food source of a given order becomes energetically viable (see Supplement S4). The resulting expression is

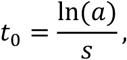

where

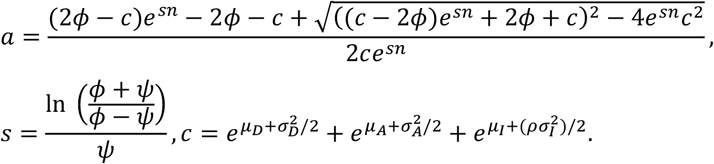

To illustrate the broader applicability of the model beyond the cannibalism framework, we include several proof-of-concept analyses involving alternative food resources in Supplement S7.

### 2.2. Application of the Model to Human Cannibalism and Justification of the Employed Parameter Values

We evaluate the expected energetic balance of cannibalism across a two-dimensional parameter space defined by trophic chain length (*ρ*) and background caloric intake (*x*). Analyses are conducted under four scenarios formed by the factorial combination of two levels of expected acquisition costs 𝔼[*c*_*A*_] and two baseline pathogen load conditions (*μ*_*I*_, *σ*_*I*_). All remaining parameters are held constant (see below for justification of exemplar values).

The specific parameter values employed are intended to illustrate the qualitative behaviour of the model rather than to provide precise empirical estimates. As such, they do not constrain the generality of the results, which follow directly from the structural properties of the model.

#### 2.2.1. Saturation Function Parameters

According to the Harris–Benedict equation

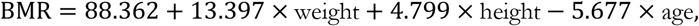

the basal metabolic rate of a 30-year-old male weighing 80 kg and measuring 180 cm in height is approximately 1850 kcal/day (Roza & Shizgal, 1984). Under conditions of very high physical activity (e.g., an active lifestyle combined with physically demanding work), daily caloric requirements are estimated at approximately 1.9 × BMR (Diabetes.co.uk, 2019), corresponding to roughly 3500 kcal/day. We adopt this value as the daily upper limit of effective caloric intake, corresponding to the asymptote of the saturation function *ϕ*.

The recommended daily caloric intake for adult men is approximately 2500 kcal/day (NHS, 2019). We use this value to parameterise the scaling constant *ψ*. While this choice implies that the marginal effectiveness of the first calories consumed may exceed 100%, this artefact has negligible influence on model behaviour; using *ψ* = 1 yields a qualitatively similar saturation function (see Supplement S3 for a direct comparison).

Model calculations are conducted on a monthly rather than daily timescale. This choice reflects the physiological constraint that humans can survive for approximately one month without food, allowing the energetic balance to be interpreted over a biologically meaningful interval. Under this framing, a cumulative balance below zero corresponds to mortality, whereas short-term deficits can be compensated by intake on other days within the same period. Surplus resources may likewise be stored, provided appropriate processing.

All monthly expressions can be reinterpreted in terms of average daily caloric intake by dividing kilocalorie values by 30, preserving interpretability. For computational consistency, we therefore rescale the saturation parameters by a factor of 30, yielding *ϕ* = 105,000 and *ψ* = 75,000 for use in the analyses that follow.

#### 2.2.2. Caloric Value of the Human Body

The total caloric value of the human body is estimated at *n* = 32,376 kcal, corresponding to 64.75 portions of 500 kcal each (Cole, 2017). The number of portions contained in a single catch is used to approximate both digestion and acquisition costs in the model.

#### 2.2.3. Digestion Costs

Baseline metabolic expenditure during chewing has been estimated at approximately 11 kcal per hour (Levine et al., 1999). We use this value as an approximation of the median digestion cost associated with processing a single portion of human meat. Although digestion costs are not limited to mastication alone, chewing provides a convenient and conservative proxy for order-of-magnitude estimation. Given the purpose of the model, moderate imprecision at this scale does not materially affect the qualitative behaviour of the results

Stochastic variation in digestion costs across portions is modelled as a multiplicative process. Relative deviations from the median are assumed to follow a log-normal distribution *e*^𝒩(0,0.1)^. Under this specification, 68.3% of portions incur digestion costs between 11*e*^−0.1^ = 9.95 and 11*e*^0.1^ = 12.16 kcal, and 95.4% fall between 11*e*^−0.2^ = 9.01 and 11*e*^0.2^ = 13.44 kcal. The expected value of this log-normal distribution is 𝔼[*c*_*Dr*_] = 11.06 kcal per portion, with a corresponding standard deviation of 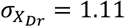. Aggregated across 64.75 portions, the total expected digestion cost is therefore 𝔼[*c*_*D*_] = 715.84 kcal (see Supplement S5 and S6 for details on distributions arising from combinations of multiplicative and additive processes).

#### 2.2.4. Acquisition Costs

We consider two acquisition cost regimes: a high-cost scenario approximating the energetic costs associated with hunting humans, and a low-cost scenario limited to food preparation.

##### 2.2.4.1. High Acquisition Cost Scenario

Even in the absence of infection risks, acquiring human meat through hunting entails substantial energetic costs, as humans constitute dangerous prey capable of inflicting serious injury. To approximate acquisition costs under this scenario, we use a physiological reference point corresponding to loss of functional capacity due to injury.

Specifically, we parameterise acquisition costs associated with hunting as a log-normal distribution with both the arithmetic mean and standard deviation set to the maximum effective caloric intake per week (3500 × 7 kcal), corresponding to 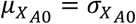 and yielding parameters *μ*_*A*0_ = 9.76 and *σ*_*A*0_ = 0.83. Under this specification, the probability of incurring costs exceeding 3500 × 7 kcal—corresponding to approximately one week of incapacitation—is approximately one third, while the probability of costs exceeding 3500 × 30 kcal—corresponding to mortality—is between one and two percent (see Supplement S8 for details based on the cumulative log-normal distribution).

Even when hunting is successful, additional energetic costs associated with food preparation are incurred. Cooking a meal for one hour has been estimated to require approximately 160 kcal (Ainsworth et al., 2011), while cooking over a fire a single meal can take approximately 30 minutes (Krebs, 2024). Based on these estimates, we set the median preparation cost of a single portion to 80 kcal.

Stochastic variation in preparation costs is modelled using a log-normal distribution with median 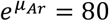 and arithmetic standard deviation 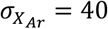, corresponding to *σ*_*Ar*_ = 0.43 (see Supplement S5). This distribution reflects the asymmetry of preparation costs: while lower costs are bounded, rare but costly events substantially increase the expected value. The resulting arithmetic mean is 𝔼[*c*_*Ar*_] = 87.89 kcal per portion.

Total expected acquisition costs correspond to the sum of expected hunting costs and the preparation of 64.75 portions. Under this scenario, total acquisition costs amount to 𝔼[*c*_*A*_] = 30,191.36 kcal, which is comparable to the total caloric value of the human body itself (Figure 3A; see Supplement S6 for details).

**Figure 3.**
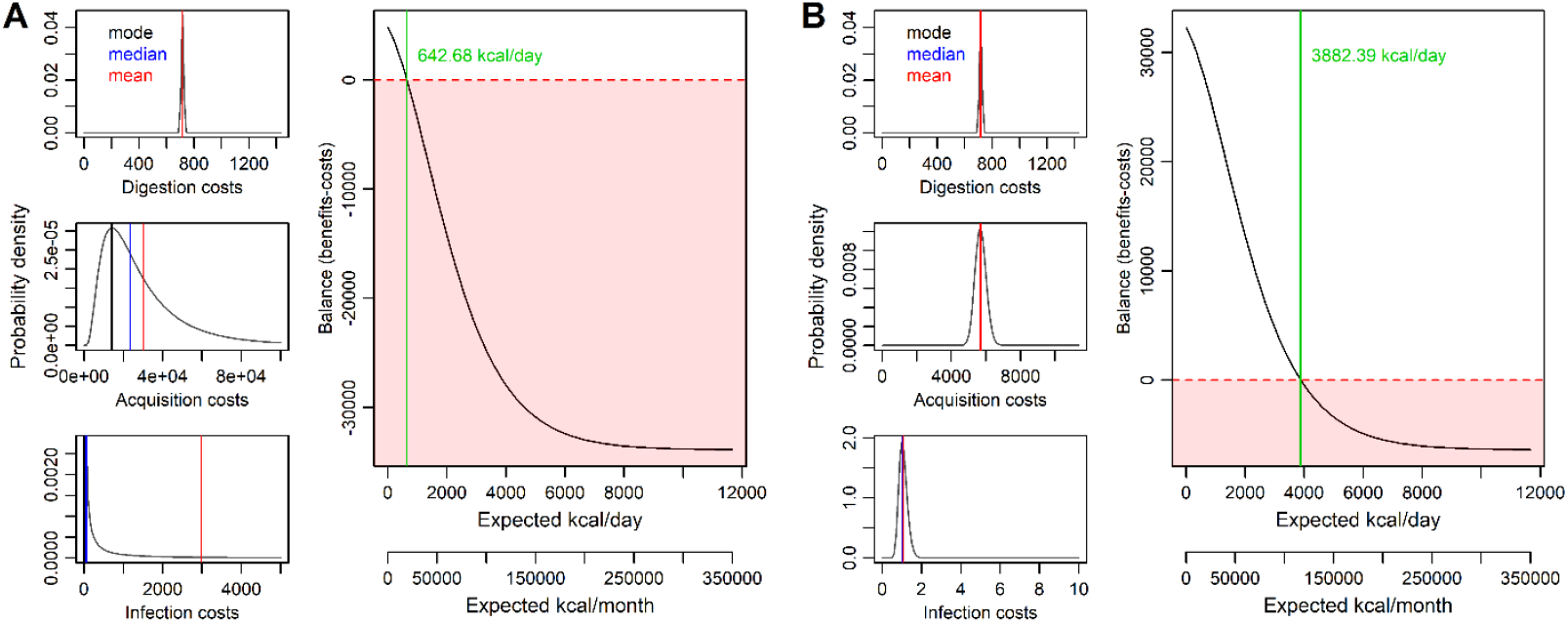
Expected energetic balance of cannibalism under two contrasting parameter regimes. *(A) High acquisition and high infection cost scenario, corresponding to raw consumption, active hunting, and prior cannibalism of the victim. Positive energetic balance occurs only under conditions of extreme food scarcity (expected caloric intake < 643 kcal/day).* *(B) Low acquisition and low infection cost scenario, corresponding to heat-processed meat, absence of prior cannibalism by the victim, and acquisition costs limited to food preparation. Under this regime, positive energetic balance persists even at comparatively high levels of background caloric intake (expected caloric intake < 3,882 kcal/day).* *Note*. Curves depict expected energetic balance as a function of background caloric intake. Insets show probability density functions of digestion, acquisition, and infection costs, with mode, median, and mean indicated. Calculations are performed on a monthly timescale.

As illustrated in Supplement S7 (Figure S7), under these acquisition costs, alternative low-risk food sources yield a higher expected energetic balance than cannibalism across a wide range of background caloric intake.

##### 2.2.4.1. Low Acquisition Cost Scenario

In the low acquisition cost scenario, hunting-related costs are omitted. This approximation corresponds to contexts in which cannibalism, for example, takes the form of ritualistic endocannibalism, involving consumption of deceased family members who die of natural causes, such as old age. Under these conditions, no additional energetic expenditure is required to obtain the body, and acquisition costs are limited to food preparation alone.

Accordingly, we use the preparation cost distribution described above for 64.75 portions, yielding total expected acquisition costs of 𝔼[*c*_*A*_] = 5,691.36 kcal. Under these conditions, the model predicts that cannibalism remains energetically viable until expected caloric intake from alternative food sources exceeds 116,479 kcal per month (equivalent to 3,882.63 kcal/day; Figure 3B).

#### 2.2.5. Infection Costs

Infection costs capture energetic losses associated with disease acquisition and immune response following consumption. While such costs are not unique to cannibalism, they play a decisive role in its energetic viability. We model infection costs using a log-normal distribution with a relatively small median (*μ*_*I*_ = 4, corresponding to *M*_*I*_ = *e*^4^) and a large standard deviation (*σ*_*I*_ = 2). Under this specification, most probability mass is concentrated at low-cost values, while the heavy right tail reflects rare but severe outcomes.

##### 2.2.5.1. High Infection Cost Scenario

The model implies that approximately one case in 1,000 results in infection costs equivalent to one week of incapacitation (*c*_*I*_ = 7 × 3500 kcal), and approximately 7 cases per 100,000 result in mortality (*c*_*I*_ = 30 × 3500 kcal). These frequencies are consistent with the assumption that severe infections are rare but non-negligible events. The expected infection cost for first-order cannibalism under this parameterisation is 𝔼[*c*_*I*_] = 403.4 kcal. In the model, infection costs are incurred once per catch, in contrast to digestion and preparation costs, which scale with the number of portions.

As the order of cannibalism increases, the variance of the infection cost distribution increases while the median remains unchanged, resulting in rapidly increasing expected infection costs. For example, consuming a first-order cannibal (i.e., second-order cannibalism) yields expected infection costs of 𝔼[*c*_*I*_] = 2,981 kcal (*μ*_*I*_ = 4, *σ*_*I*_ = 2, *ρ* = 2). Expected infection costs associated with third-order cannibalism exceed the total caloric value of the human body (𝔼[*c*_*I*_] = 162,754.8 kcal), rendering such behaviour energetically unviable under this regime.

##### 2.2.5.2. Low Infection Costs Scenario

Humans can reduce pathogen load through food processing, most notably using fire. Thermal processing substantially reduces the viability of many pathogens; accordingly, in the low infection cost scenario we assume a hundredfold reduction in baseline pathogen load, yielding distribution parameters *μ*_*I*_ = 0.04 and *σ*_*I*_ = 0.2. As discussed above, the distribution’s variance, not standard deviation, scales linearly with the factor of multiplication or division.

Under this parameterisation, expected infection costs associated with third-order cannibalism remain low (𝔼[*c*_*I*_] = 1.13 kcal), whereas at sufficiently high orders of cannibalism infection costs again exceed energetic benefits (e.g., 𝔼[*c*_*I*_] = 32,859.63 kcal for *ρ* = 517). Even substantial reductions in baseline pathogen load cannot eliminate the exponential scaling of infection costs with trophic chain length.

Residual infection risk is attributed to durable pathogens that are resistant to standard heat-based processing. Prion diseases, for instance, are rare but prions (misfolded proteins that induces folding problems in normal variants of identical amino acid chains) are resistant to denaturation by heat typically sufficient to eliminate viruses and bacteria (Aguzzi & Heppner, 2000).

## 3. Results

Model outcomes are visualised in Figures 3 and 4. Across all parameterisations, expected energetic balance depends jointly on background caloric intake, acquisition costs, infection costs, and the order of cannibalism.

**Figure 4.**
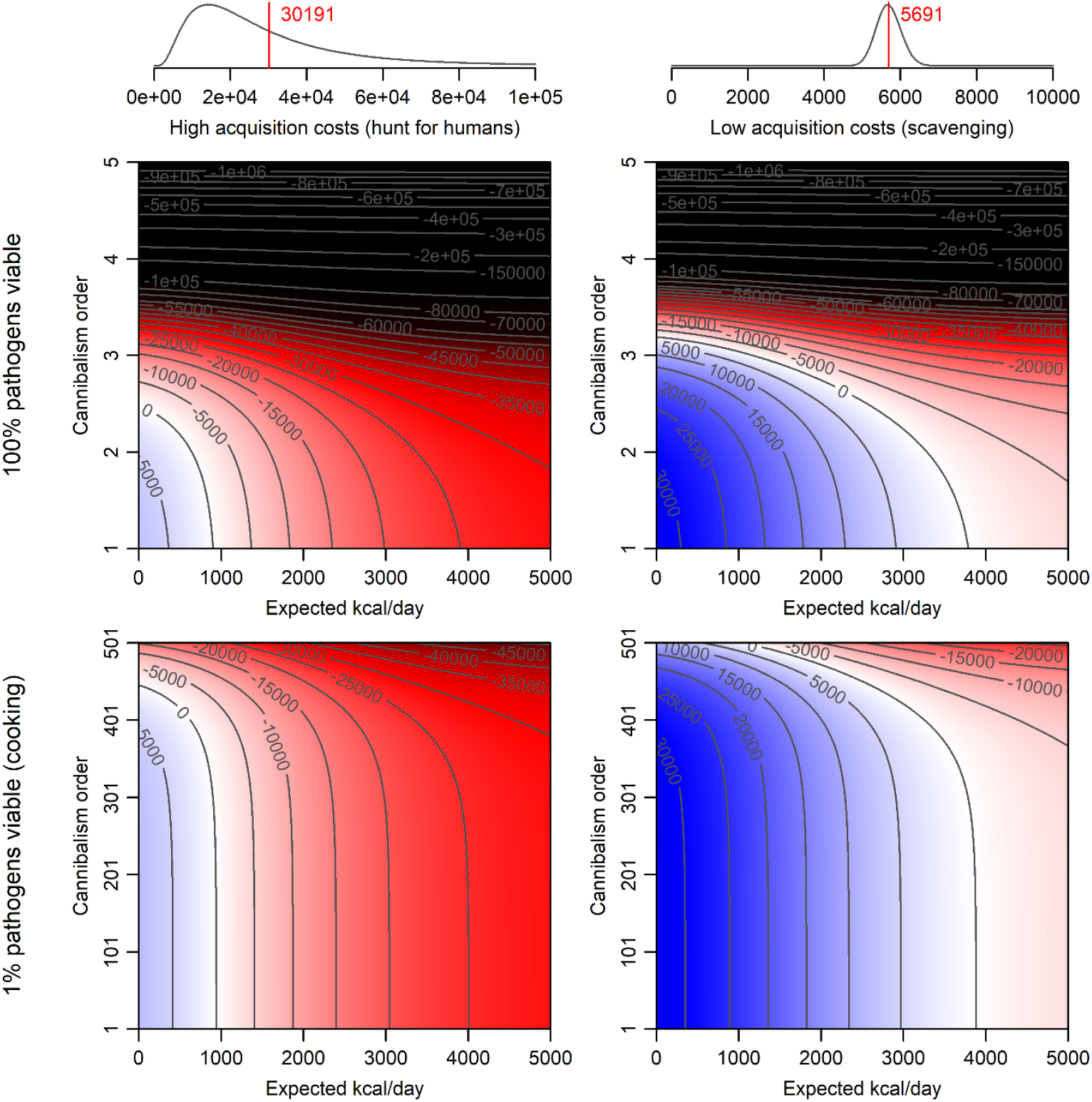
Expected energetic balance of human body consumption as a function of food abundance and cannibalism order under varying acquisition and infection cost regimes. *Note*. Contour plots show expected energetic balance across background caloric intake (x-axis) and cannibalism order (y-axis). Columns correspond to high acquisition costs (hunting humans) and low acquisition costs (scavenging), respectively. Rows correspond to high infection costs (100% pathogen viability; raw consumption) and low infection costs (1% pathogen viability; heat-processed meat). Red tones indicate negative expected balance, whereas blue tones indicate positive expected balance. Digestion costs are held constant at 11.05 kcal per 500 kcal portion of human meat. Profitability calculations are performed on a monthly timescale, with saturation function scaling parameters *ϕ*=30 × 3500 and *n*=32,376 kcal. Infection costs for 100% pathogen viability are characterised by a log-normal distribution with *μ*_*I*_ = 4 and *σ*_*I*_ = 2. Note the difference in scale on the vertical axis between the upper and lower rows.

First, cannibalism is energetically viable primarily under conditions of low acquisition costs. Scenarios in which human flesh is obtained without hunting (e.g., as a by-product of other processes) yield substantially larger regions of positive energetic balance than scenarios requiring active pursuit of human prey (Figures 3 and 4; right vs left columns).

Second, the energetic viability of cannibalism is strongly constrained by the order of cannibalism. Positive expected balance is restricted to low trophic chain lengths when baseline infection costs are high, with the region of profitability rapidly shrinking as cannibalism order increases (Figure 4, upper panels). This constraint is relaxed under reduced pathogen load, allowing positive balance to persist across substantially higher cannibalism orders (Figure 4, lower panels).

Third, food processing substantially alters the shape of the profitable region. Under conditions of reduced infection costs associated with heat processing, the parameter space yielding positive energetic balance expands both along the background caloric intake axis and the cannibalism order axis (Figure 4, bottom row). In contrast, when meat is consumed raw, profitable cannibalism is limited to a narrow range of low-order chains and low background caloric intake (Figure 4, top row).

Fourth, profitable cannibalism emerges only under sufficiently low background caloric intake when acquisition costs are high, even if cannibalism order is minimal (Figure 3A). Under low acquisition and low infection cost regimes, however, positive energetic balance can be maintained even at comparatively high levels of background caloric intake (Figure 3B).

Finally, infection costs scale differently from acquisition costs in cooperative contexts (see Supplement S7 for further discussion, Figure S11 for an example). Whereas acquisition costs are divided among cooperating individuals, infection costs are borne individually. As a consequence, prey types associated with higher expected infection costs yield smaller regions of positive energetic balance under shared consumption compared to prey with lower expected infection costs, holding total energetic intake constant (Figure 4).

Supplementary analyses further indicate that isolated cannibalistic events under extreme conditions yield positive energetic balance across a wide range of cannibalism orders (Supplement S8).

## 4. Discussion

This project set out to address a long-standing puzzle in the study of human cannibalism: why has cannibalism emerged repeatedly across human history, yet remained strongly tabooed in both historical and contemporary societies? We approached this question by assuming that cannibalism involves a trade-off, in which specific ecological conditions can make the practice temporarily viable, while accumulating costs eventually suppress it. To formalise this idea, we modelled cannibalism as a balance between nutritional benefits and multiple sources of cost, with particular attention to infection risks that increase as cannibalism is repeated across individuals. The results show that cannibalism can yield a positive energetic balance under a narrow range of conditions, such as severe food scarcity, low acquisition costs, and reduced infection risk through food processing (e.g., the use of fire). However, as cannibalism extends across longer chains of consumers, expected infection costs rise sharply and rapidly outweigh any nutritional gains.

### 4.1. Interpretative Implications

The present model is not intended as a purely abstract or speculative exercise. Instead, it provides a structured way of thinking about the conditions under which cannibalism is more or less likely to occur, and about the constraints that should limit its form and persistence. The model provides a point of reference for archaeological and anthropological interpretation, helping to assess which ecological and behavioural explanations of cannibalistic evidence are probable.

A central interpretative implication of the model is that cannibalism is inherently high-risk and therefore unlikely to be practiced regularly over long periods. This suggests that claims of widespread or habitual cannibalism should be treated with caution, particularly when such practices are inferred during periods of relative abundance rather than starvation (Bello et al., 2015; Carbonell et al., 2010; Marginedas et al., 2025). At the same time, this perspective aligns with ethnographic critiques arguing that cannibalism has often been inferred too readily in accounts of other populations (Pickering, 2017). Even when cannibalism is interpreted as a survival strategy, the model implies that it could only operate temporarily, under conditions of food scarcity, and within narrow limits, consistent with views that treat cannibalism as a latent human psychological disposition regulated by cultural norms and activated primarily during ecological stress (Petrinovich, 2000).

Acquisition costs are critical for energetic viability. Cannibalism, therefore, is most likely to take the form of endocannibalism, for example through the consumption of deceased group members. Exocannibalism, by contrast, involves substantially higher acquisition costs, as humans represent dangerous prey. Even where evidence points to exocannibalism, the model suggests that cannibalism is more plausibly interpreted as a by-product of warfare or raiding rather than as a purely nutritional strategy (for example, in the case of the Māori; Moon, 2008). Exocannibalism appears to have been rare and, when practiced, organised to minimise risk, for instance by preferentially targeting women and children rather than adult males (Cosnefroy et al., 2025; Fernández-Jalvo et al., 1999; Rodríguez et al., 2019; Saladié et al., 2012). Furthermore, targeting younger individuals would be expected to reduce infection risk, because they are less likely to have participated in prior cannibalistic events, thereby limiting trophic chain length.

The model also highlights the importance of fire and food processing for the viability of cannibalism. By reducing baseline infection risk, cooking substantially expands the conditions under which cannibalism can yield a positive energetic balance, making it a more plausible option for hominins following the control of fire (ca. 400 ka), potentially beginning with *Homo erectus* (Wrangham, 2017). Conversely, the model implies that regular cannibalism would have been far less viable for hominin populations lacking food-processing technologies, as infection costs would escalate rapidly even under severe ecological stress. Accordingly, assemblages displaying burning or other signs of cooking provide stronger support for cannibalism interpretations than accounts emphasising raw consumption, which imply higher infection costs and lower plausibility outside short-term crises (Marlar et al., 2000; Saladié et al., 2025; Vilaca, 2000). Even with cooking, however, infection costs increase nonlinearly with repeated within-species consumption, limiting cannibalism to episodic rather than sustained practice.

Taken together, the results imply that if cannibalism was practiced over a prolonged period, it must have occurred under a highly restricted set of conditions: during severe food scarcity, through the consumption of individuals whose bodies could be obtained at minimal acquisition risk, and with the use of fire or other food-processing techniques. Given the structure of the model, this configuration is, mathematically speaking, the only one compatible with sustained energetic viability and should therefore serve as the default interpretative assumption. Claims that cannibalism functioned as a stable subsistence strategy outside periods of starvation, involved systematic exocannibalism, or relied on the consumption of raw human flesh thus require explicit supporting evidence. In the absence of such evidence, these higher-risk interpretations are less consistent with the constraints imposed by the model.

### 4.2. Cultural Evolution of Cannibalism

The model shows that cannibalism is not a viable long-term subsistence strategy. It also implies that cannibalistic behaviour need not be immediately eliminated by natural selection. Most cannibalistic episodes do not result in the immediate death of the cannibal, even at higher cannibalism orders, allowing populations time to respond before catastrophic collapse. At the same time, the model predicts that costs escalate nonlinearly with repeated within-species consumption, such that unrestricted cannibalism inevitably becomes unsustainable unless cannibalism order is actively constrained. This structural property helps explain why sustained cannibalism is rare in humans and why, in non-human animals, cannibalism is most often restricted to the consumption of newborn or very young individuals (Fox, 1975; Polis, 1981). In animals, this restraint may arise through natural selection acting on aversions to adult conspecifics, particularly in the absence of food-processing technologies that reduce pathogen viability (see Figure 4), whereas in humans it is more effectively achieved through cultural regulation (Chudek & Henrich, 2011).

From this perspective, cannibalism taboos can be understood as adaptive cultural responses to escalating epidemiological risks (Pooladvand et al., 2024). Rather than functioning solely as absolute prohibitions, taboos operate as regulatory systems that specify who may be consumed, under what conditions, and how often (Hong, 2024). By constraining cannibalism order, through age-based restrictions, distinctions between endo- and exocannibalism, or tightly circumscribed contexts, taboos reduce long-term infection risk while still allowing limited departures under extreme ecological stress. Cultural systems that failed to regulate cannibalism through taboos would therefore be expected to disappear.

A related but distinct issue concerns the role of ritual in the emergence of cannibalistic practices. Because the model shows that cannibalism entails substantial and escalating costs, it provides little support for treating ritual cannibalism as a distinct category characterised by high costs and no clear benefits. Sharp distinctions between “ritual,” and “survival” cannibalism are therefore conceptually misleading (Bello, 2024; Byard, 2023). Ritual is better understood not as an alternative form of cannibalism, but as a feature that commonly accompanies other forms, enabling participation and regulating practice under conditions in which cannibalism becomes temporarily viable.

Moving from a state of complete prohibition to the consumption of human flesh requires overcoming strong emotional and moral barriers (Haidt et al., 2000; Horberg et al., 2009; Misiak et al., 2025). Ritualisation can facilitate this transition by framing cannibalism as a sanctioned, meaningful, and socially coordinated act, thereby reducing individual resistance to consuming an otherwise prohibited food. Psychological research shows that rituals can enable the consumption of unusual or morally challenging foods (Carter et al., 2025), and human flesh represents an extreme case of such resistance. Rituals may also contribute to the maintenance of these norms once established (Chvaja et al., 2023; Rossano, 2012). This interpretation aligns with ethnographic evidence indicating that cannibalism is often embedded within ritual frameworks, regardless of whether it occurs in subsistence, mortuary, or conflict-related contexts (Lindenbaum, 2004; Moon, 2008; Schutt, 2018).

Because the use of fire limits the baseline pool of pathogens capable of adapting to trophic transmission, cannibalism among humans should, all else equal, occur more frequently than cannibalism among other animals. Irrespective of narratives that frame it primarily as a tool of outgroup dehumanization (Arens, 1979; Pickering, 2017). Repeated episodes of ritualised, transgenerational, cooking-facilitated cannibalism could create selective environments favouring increased resistance to prion diseases, as suggested by genetic evidence (Mead et al., 2003). In the model, a hundredfold reduction in pathogen load substantially (even more than hundredfold) expands the range of cannibalism orders under which consumption yields a positive energetic balance. If supported by further empirical evidence, this pattern would constitute a clear example of gene–culture coevolution (Boyd & Richerson, 2024; Feldman & Laland, 1996).

However, even under these circumstances, a taboo that ultimately prohibits or drastically restricts cannibalism must emerge if a society is to survive. No process of biological adaptation, assumed to be linear in a population of fixed size under additive genetics (Tureček et al., 2019), can keep pace with the exponential growth of expected infection costs. This contrast highlights the greater degree of manoeuvring space available to humans relative to other animals. In non-human species, where high-fidelity transgenerational social transmission is absent, strict aversion to the consumption of adult conspecifics represents the only plausible mechanism preventing the spread of specialised trophic pathogens at moderate cannibalism orders. Human societies, by contrast, can construct and reliably transmit norms that permit cannibalism only under conditions that occur rarely, for example once every few generations. When sequences of cannibals consuming cannibals are regularly interrupted, conspecific meat can constitute a viable, but tightly constrained, dietary extension.

### 4.3. Limitations and Future Directions

The present model deliberately adopts a simplified representation of nutritional benefits by treating food value exclusively in terms of kilocalories. In reality, nutrition is much more complicated. Mixed evidence suggests that cannibalism may, in some contexts, be associated with deficiencies in specific macro- or micronutrients rather than with absolute caloric scarcity (Guil-Guerrero, 2017; Ortiz de Montellano, 1978). Nutrient-driven motivations could, in principle, render cannibalism viable even when energetic requirements are otherwise met. Such considerations are not captured by the current formulation and represent an important direction for future extensions of the model, particularly in contexts where protein, fatty acids, or micronutrients may have been limiting.

A second limitation concerns the structure of trophic chains. In the present model, cannibalism order (ρ) is treated as a linear chain in which each individual consumes one other cannibal. In reality, a single individual could consume multiple cannibals, potentially amplifying infection risk. Extending the model to include a parameter θ representing the number of cannibals consumed by a single individual would require incorporating extreme-value processes, for example through Gumbel or related generalised extreme value distributions. Under such an extension, the infection risk posed by a cannibal of order ρ would reflect the distribution of maxima drawn from multiple lower-order chains rather than a single lineage. While this added complexity could refine estimates of infection costs, the present parameter ρ can be interpreted, with minimal loss of generality, as the expected longest trophic chain in an individual’s diet, which dominates the expected value of infection costs. As such, the current formulation captures the core scaling logic while remaining analytically tractable.

More generally, the model relies on illustrative parameter values and functional forms rather than precise empirical estimates. These choices are intended to demonstrate the qualitative behaviour of the system rather than to provide exact quantitative predictions. We therefore welcome further discussion regarding the most appropriate parameterisations and functional assumptions. Nonetheless, we are confident that the initial estimates used here fall within a plausible biological range and that the central results follow from the structural properties of the model rather than from fine-tuned parameter choices.

Finally, the present analysis treats cannibalism primarily as a subsistence strategy evaluated in terms of energetic balance. This focus necessarily excludes potential social benefits that could, under some conditions, offset costs. Cannibalism may function as a costly signal of formidability, hunting ability, or unpredictability, or as a means of terrorising rival groups by cultivating a reputation for extreme behaviour (Fessler et al., 2014; Kleef et al., 2023). Macabre accounts of indigenous cannibalism may be inflated by Eurocentric xenophobia, but an equally important source of exaggeration may stem from active efforts to impress potentially hostile outgroups. Formalising these social payoffs would require a different modelling framework, potentially incorporate signalling theory or intergroup competition, and lie beyond the scope of the present study. Exploring how such individual-level benefits interact with the population-level costs identified here represents a promising direction for future research.

### 4.4. Conclusions

This paper addresses why cannibalism has repeatedly appeared in human history while remaining one of the strongest and most persistent cultural taboos. By formalising cannibalism as a trade-off between nutritional benefits and escalating costs (most critically, infection risks that increase nonlinearly with repeated within-species consumption) we show that cannibalism is viable only under narrow and highly constrained conditions. These include extreme resource scarcity, low acquisition costs, reduced pathogen load through food processing, and strict limits on cannibalism order. Outside this parameter space, costs rapidly outweigh benefits, rendering sustained cannibalism unviable. This structure accounts for the episodic appearance of cannibalism in the archaeological and ethnographic record and its consistent regulation or suppression through cultural norms. Cannibalism taboos thus emerge not as arbitrary moral prohibitions, but as predictable cultural responses to epidemiological constraints. More broadly, the model illustrates how biological limits on disease transmission can shape cultural evolution by favouring regulatory systems that permit rare exceptions under extreme conditions while preventing long-term population collapse.

## References

Aguzzi, A., & Heppner, F. L. (2000). Pathogenesis of prion diseases: A progress report. Cell Death and Differentiation, 7(10), 889–902.

Ainsworth, B. E., Haskell, W. L., Herrmann, S. D., Meckes, N., Bassett, D. R., Tudor-Locke, C., Greer, J. L., Vezina, J., Whitt-Glover, M. C., & Leon, A. S. (2011). 2011 Compendium of Physical Activities: A Second Update of Codes and MET Values. Medicine & Science in Sports & Exercise, 43(8), 1575–1581. 10.1249/MSS.0b013e31821ece12

Alpers, M. P. (2008). The epidemiology of kuru: Monitoring the epidemic from its peak to its end. Philosophical Transactions of the Royal Society B: Biological Sciences, 363(1510), 3707–3713. 10.1098/rstb.2008.0071

Arens, W. (1979). The Man-Eating Myth: Anthropology and Anthropophagy: Anthropology and Anthropophagy. Oxford University Press, USA.

Bello, S. M. (2024). The Archaeology of Cannibalism: A Review of the Taphonomic Traits Associated with Survival and Ritualistic Cannibalism. Journal of Archaeological Method and Theory, 32(1), 11. 10.1007/s10816-024-09676-3

Bello, S. M., Saladié, P., Cáceres, I., Rodríguez-Hidalgo, A., & Parfitt, S. A. (2015). Upper Palaeolithic ritualistic cannibalism at Gough’s Cave (Somerset, UK): The human remains from head to toe. Journal of Human Evolution, 82, 170–189. 10.1016/j.jhevol.2015.02.016

Boyd, R., & Richerson, P. J. (2024). Cultural evolution: Where we have been and where we are going (maybe). Proceedings of the National Academy of Sciences, 121(48), e2322879121. 10.1073/pnas.2322879121

Byard, R. W. (2023). Cannibalism—Overview and medicolegal issues. Forensic Science, Medicine and Pathology, 19(2), 281–287. 10.1007/s12024-023-00623-4

Carbonell, E., Cáceres, I., Lozano, M., Saladié, P., Rosell, J., Lorenzo, C., Vallverdú, J., Huguet, R., Canals, A., & Bermúdez de Castro, J. M. (2010). Cultural Cannibalism as a Paleoeconomic System in the European Lower Pleistocene. Current Anthropology, 51(4), 539–549. 10.1086/653807

Carter, A. G., Faber, N. S., & McGuire, L. (2025). Values Over Virtues: How Children Trade Off Their Moral Concern for Animals With the Importance of Human Eating Practices. Social Psychological and Personality Science, 19485506251398890.

Chudek, M., & Henrich, J. (2011). Culture–gene coevolution, norm-psychology and the emergence of human prosociality. Trends in Cognitive Sciences, 15(5), 218–226. 10.1016/j.tics.2011.03.003

Chvaja, R., Horský, J., Lang, M., & Kundt, R. (2023). Positive association between ritual performance and perceived objectivity of moral norms. The International Journal for the Psychology of Religion, 33(2), 115–135.

Claessen, D., de Roos, A. M., & Persson, L. (2004). Population dynamic theory of size–dependent cannibalism. Proceedings of the Royal Society B: Biological Sciences, 271(1537), 333–340. 10.1098/rspb.2003.2555

Cole, J. (2017). Assessing the calorific significance of episodes of human cannibalism in the Palaeolithic. Scientific Reports, 7(1), 44707. 10.1038/srep44707

Cosnefroy, Q., Crevecoeur, I., Semal, P., Hajdinjak, M., Bossoms Mesa, A., Krause, J., Gnecchi-Ruscone, G. A., Posth, C., Bocherens, H., Devièse, T., & Rougier, H. (2025). Highly selective cannibalism in the Late Pleistocene of Northern Europe reveals Neandertals were targeted prey. Scientific Reports, 15(1), 40741. 10.1038/s41598-025-24460-3

Diabetes.co.uk. (2019). BMR Calculator. Diabetes.Co.Uk. https://www.diabetes.co.uk/bmr-calculator.html

Feldman, M. W., & Laland, K. N. (1996). Gene-culture coevolutionary theory. Trends in Ecology & Evolution, 11(11), 453–457. 10.1016/0169-5347(96)10052-5

Fernández-Jalvo, Y., Carlos Diez, J., Cáceres, I., & Rosell, J. (1999). Human cannibalism in the Early Pleistocene of Europe (Gran Dolina, Sierra de Atapuerca, Burgos, Spain). Journal of Human Evolution, 37(3), 591–622. 10.1006/jhev.1999.0324

Fessler, D. M. T., Tiokhin, L. B., Holbrook, C., Gervais, M. M., & Snyder, J. K. (2014). Foundations of the Crazy Bastard Hypothesis: Nonviolent physical risk-taking enhances conceptualized formidability. Evolution and Human Behavior, 35(1), 26–33. 10.1016/j.evolhumbehav.2013.09.003

Fisher, R. A. (1918). XV.—The Correlation between Relatives on the Supposition of Mendelian Inheritance. Earth and Environmental Science Transactions of The Royal Society of Edinburgh, 52(2), 399–433. 10.1017/S0080456800012163

Fox, L. R. (1975). Cannibalism in Natural Populations. Annual Review of Ecology and Systematics, 6, 87–106.

Guil-Guerrero, J. L. (2017). Evidence for chronic omega-3 fatty acids and ascorbic acid deficiency in Palaeolithic hominins in Europe at the emergence of cannibalism. Quaternary Science Reviews, 157, 176–187. 10.1016/j.quascirev.2016.12.016

Haidt, J., Bjorklund, F., & Murphy, S. (2000). Moral dumbfounding: When intuition finds no reason. Unpublished Manuscript, University of Virginia, 191, 221.

Horberg, E. J., Oveis, C., Keltner, D., & Cohen, A. B. (2009). Disgust and the moralization of purity. Journal of Personality and Social Psychology, 97(6), 963–976. 10.1037/a0017423

Kleef, G. A. van, Wanders, F., Vianen, A. E. M. van, Dunham, R. L., Du, X., & Homan, A. C. (2023). Rebels with a cause? How norm violations shape dominance, prestige, and influence granting. PLOS ONE, 18(11), e0294019. 10.1371/journal.pone.0294019

Klitzman, R. L., Alpers, M. P., & Gajdusek, D. C. (1985). The Natural Incubation Period of Kuru and the Episodes of Transmission in Three Clusters of Patients. Neuroepidemiology, 3(1), 3–20. 10.1159/000110837

Kraft, T. S., Venkataraman, V. V., Wallace, I. J., Crittenden, A. N., Holowka, N. B., Stieglitz, J., Harris, J., Raichlen, D. A., Wood, B., Gurven, M., & Pontzer, H. (2021). The energetics of uniquely human subsistence strategies. Science, 374(6575), eabf0130. 10.1126/science.abf0130

Krebs, J. (2024, April 25). How to Cook Over a Fire, According to a Survival Instructor. Backpacker. https://www.backpacker.com/survival/how-to-cook-over-a-fire/

Lafferty, K. D. (1999). The Evolution of Trophic Transmission. Parasitology Today, 15(3), 111–115. 10.1016/S0169-4758(99)01397-6

Levine, J., Baukol, P., & Pavlidis, I. (1999, December 30). The Energy Expended in Chewing Gum [Letter]. Massachusetts Medical Society. (world). 10.1056/NEJM199912303412718

Liberski, P. P., Sikorska, B., Lindenbaum, S., Goldfarb, L. G., McLean, C., Hainfellner, J. A., & Brown, P. (2012). Kuru: Genes, Cannibals and Neuropathology. Journal of Neuropathology & Experimental Neurology, 71(2), 92–103. 10.1097/NEN.0b013e3182444efd

Lindenbaum, S. (2004). Thinking About Cannibalism. Annual Review of Anthropology, 33(Volume 33, 2004), 475–498. 10.1146/annurev.anthro.33.070203.143758

Macbeth, H., Schiefenhövel, W., & Collinson, P. (2007). Cannibalism No Myth, But Why So Rare? In Consuming the inedible: Neglected dimensions of food choice (Vol.6, p. 189). Berghahn Books. https://www.google.com/books?hl=pl&lr=&id=AgNoh5r0kuoC&oi=fnd&pg=PA189&dq=CANNIBALISM+NO+MYTH,+BUT+WHY+SO+RARE%3F&ots=XrdBKd2ilt&sig=9ObImQYLDTG6i5yxvmOXYnee8FE

Marginedas, F., Saladié, P., Połtowicz-Bobak, M., Terberger, T., Bobak, D., & Rodríguez-Hidalgo, A. (2025). New insights of cultural cannibalism amongst Magdalenian groups at Maszycka Cave, Poland. Scientific Reports, 15(1), 2351. 10.1038/s41598-025-86093-w

Marlar, R. A., Leonard, B. L., Billman, B. R., Lambert, P. M., & Marlar, J. E. (2000). Biochemical evidence of cannibalism at a prehistoric Puebloan site in southwestern Colorado. Nature, 407(6800), 74–78. 10.1038/35024064

Mathews, J., Glasse, R., & Lindenbaum, S. (1968). KURU AND CANNIBALISM. The Lancet, Originally Published as Volume 2, Issue 7565, 292(7565), 449–452. 10.1016/S0140-6736(68)90482-0

McHugh, C., McGann, M., Igou, E. R., & Kinsella, E. L. (2017). Searching for Moral Dumbfounding: Identifying Measurable Indicators of Moral Dumbfounding. Collabra: Psychology, 3(1), 23. 10.1525/collabra.79

Mead, S., Stumpf, M. P. H., Whitfield, J., Beck, J. A., Poulter, M., Campbell, T., Uphill, J. B., Goldstein, D., Alpers, M., Fisher, E. M. C., & Collinge, J. (2003). Balancing Selection at the Prion Protein Gene Consistent with Prehistoric Kurulike Epidemics. Science. (world). 10.1126/science.1083320

Misiak, M., Paruzel-Czachura, M., Butovskaya, M., Karwowski, M., & Sorokowski, P. (2025). Unearthing the Foundations: Testing the Universality of Moral Foundations Theory in Three Small-Scale Populations (9w42p_v1). OSF Preprints. 10.31219/osf.io/9w42p_v1

Moon, P. (2008). This Horrid Practice. Penguin Random House New Zealand Limited.

NHS. (2019). What should my daily intake of calories be? National Health Service. https://www.nhs.uk/common-health-questions/food-and-diet/what-should-my-daily-intake-of-calories-be/

Oldak, S. E., Maristany, A. J., & Sa, B. C. (2023). Wendigo Psychosis and Psychiatric Perspectives of Cannibalism: A Complex Interplay of Culture, Psychology, and History. Cureus, 15(10), e47962. 10.7759/cureus.47962

Ortiz de Montellano, B. R. (1978). Aztec Cannibalism: An Ecological Necessity? Science, 200(4342), 611–617. 10.1126/science.200.4342.611

Park, A. W. (2019). Food web structure selects for parasite host range. Proceedings of the Royal Society B: Biological Sciences, 286(1908), 20191277. 10.1098/rspb.2019.1277

Petrinovich, L. F. (2000). The Cannibal Within. Transaction Publishers.

Pickering, M. (2017). Cannibalism in the Ethnographic Record. In The International Encyclopedia of Anthropology (pp. 1–10). John Wiley & Sons, Ltd. 10.1002/9781118924396.wbiea1993

Polis, G. A. (1981). The Evolution and Dynamics of Intraspecific Predation. Annual Review of Ecology and Systematics, 12, 225–251.

Pooladvand, P., Kendal, J. R., & Tanaka, M. M. (2024). How cultural innovations trigger the emergence of new pathogens. Proceedings of the National Academy of Sciences, 121(48), e2322882121. 10.1073/pnas.2322882121

Pozio, E., Rinaldi, L., Marucci, G., Musella, V., Galati, F., Cringoli, G., Boireau, P., & La Rosa, G. (2009). Hosts and habitats of Trichinella spiralis and Trichinella britovi in Europe. International Journal for Parasitology, 39(1), 71–79. 10.1016/j.ijpara.2008.06.006

Rodríguez, J., Guillermo, Z.-R., & Ana, M. (2019). Does optimal foraging theory explain the behavior of the oldest human cannibals? Journal of Human Evolution, 131, 228–239. 10.1016/j.jhevol.2019.03.010

Rossano, M. J. (2012). The essential role of ritual in the transmission and reinforcement of social norms. Psychological Bulletin, 138(3), 529–549. 10.1037/a0027038

Roza, A. M., & Shizgal, H. M. (1984). The Harris Benedict energy requirements equation reevaluated: Resting and the body cell mass. The American Journal of Clinical Nutrition, 40(July), 168–182.

Rudolf, V. H. W. (2007). The interaction of cannibalism and omnivory: Consequences for community dynamics. Ecology, 88(11), 2697–2705. 10.1890/06-1266.1

Rudolf, V. H. W., & Antonovics, J. (2007). Disease transmission by cannibalism: Rare event or common occurrence? Proceedings of the Royal Society B: Biological Sciences, 274(1614), 1205–1210. 10.1098/rspb.2006.0449

Saladié, P., Cáceres, I., Huguet, R., Rodríguez-Hidalgo, A., Santander, B., Ollé, A., Gabucio, M. J., Martín, P., & Marín, J. (2015). Experimental Butchering of a Chimpanzee Carcass for Archaeological Purposes. PLOS ONE, 10(3), e0121208. 10.1371/journal.pone.0121208

Saladié, P., Huguet, R., Rodríguez-Hidalgo, A., Cáceres, I., Esteban-Nadal, M., Arsuaga, J. L., Bermúdez de Castro, J. M., & Carbonell, E. (2012). Intergroup cannibalism in the European Early Pleistocene: The range expansion and imbalance of power hypotheses. Journal of Human Evolution, 63(5), 682–695. 10.1016/j.jhevol.2012.07.004

Saladié, P., Marginedas, F., Morales, J. I., Vergès, J. M., Allué, E., Expósito, I., Lozano, M., Martín, P., Iglesias-Bexiga, J., Fontanals, M., Marsal, R., Hernando, R., Burguet-Coca, A., & Rodríguez-Hidalgo, A. (2025). Evidence of neolithic cannibalism among farming communities at El Mirador cave, Sierra de Atapuerca, Spain. Scientific Reports, 15, 26648. 10.1038/s41598-025-10266-w

Schutt, B. (2018). Cannibalism: A Perfectly Natural History. Algonquin Books.

Smith, A. R., Carmody, R. N., Dutton, R. J., & Wrangham, R. W. (2015). The significance of cooking for early hominin scavenging. Journal of Human Evolution, 84, 62–70. 10.1016/j.jhevol.2015.03.013

Sugg, R. (2015). Mummies, Cannibals and Vampires: The History of Corpse Medicine from the Renaissance to the Victorians (2nd edn). Routledge. 10.4324/9781315666365

Turecek, P., Slavík, J., Kozák, M., & Havlíček, J. (2019). Non-particulate inheritance revisited: Evolution in systems with Parental Variability-Dependent Inheritance. Biological Journal of the Linnean Society, 127(2), 518–533. 10.1093/biolinnean/blz041

Vilaca, A. (2000). Relations between Funerary Cannibalism and Warfare Cannibalism: The Question of Predation. Ethnos, 65(1), 83–106. 10.1080/001418400360652

Vilaça, A. (2005). Chronically Unstable Bodies: Reflections on Amazonian Corporalities. Journal of the Royal Anthropological Institute, 11(3), 445–464. 10.1111/j.1467-9655.2005.00245.x

Winterhalder, B., & Smith, E. A. (2000). Analyzing adaptive strategies: Human behavioral ecology at twenty-five. Evolutionary Anthropology Issues News and Reviews, 9(2), 51–72. 10.1002/(sici)1520-6505(2000)9:2%253C51::aid-evan1%253E3.0.co;2-7

Yustos, M., Terreros, J. Y. S. de los, Yustos, M., & Terreros, J. Y. S. de los. (2015). Cannibalism in the Neanderthal World: An Exhaustive Revision. Journal of Taphonomy, 13(1), 33–52.

