## Supplement for "The Cannibalistic Trade-Off: Why Human Cannibalism Emerges and Why Taboos Suppress It"

**Supplementary material**

**S1. Algebraic construction of the saturation function**

To construct a plausible saturation function, we draw inspiration in the inverse-logit function:

$$f(x)=\frac{e^{x}}{e^{x}+1},$$

which naturally occupies the function domain $[0.5,1)$ for $x>0$. We subtract $0.5$ from this formula to achieve $f(0)=0$ and multiply it by $2$ to stretch the domain from $[0,0.5)$ to $[0,1)$.

Thus the baseline saturation function is:

$$f(x)=\frac{2e^{x}}{e^{x}+1}-1$$

the $y$ scaling parameter in just $\phi$ (we multiply the expression by the constant):

$$f(x)=\frac{2\phi e^{x}}{e^{x}+1}-\phi$$

To get the $x$ scaling parameter $s$, we just solve for $s$ in

$$\frac{2\phi e^{\psi s}}{e^{\psi s}+1}-\phi=\psi.$$

Add $\phi$ to both sides:

$$\frac{2\phi e^{\psi s}}{e^{\psi s}+1}=\phi+\psi$$

Multiply both sides by $e^{\psi s}+1$:

$$2\phi e^{\psi s}=\phi e^{\psi s}+\psi e^{\psi s}+\phi+\psi$$

Subtract the elements on the right side that contain $e^{\psi s}$ from both sides:

$$2\phi e^{\psi s}-\phi e^{\psi s}-\psi e^{\psi s}=\phi+\psi$$

Factor out $e^{\psi s}$:

$$(2\phi-\phi-\psi)e^{\psi s}=\phi+\psi$$

$$(\phi-\psi)e^{\psi s}=\phi+\psi$$

Divide both sides by $\phi-\psi$:

$$e^{\psi s}=\frac{\phi+\psi}{\phi-\psi}$$

Taking the natural logarithm of both sides yields:

$$\psi s=\text{ln}\left( \frac{\phi+\psi}{\phi-\psi} \right)$$

Divide by $\psi$:

$$s=\frac{\text{ln}\left( \frac{\phi+\psi}{\phi-\psi} \right)}{\psi}$$

The general saturation function is then:

$$f(x)=\frac{2\phi e^{\frac{\text{ln}\left( \frac{\phi+\psi}{\phi-\psi} \right)x}{\psi}}}{e^{\frac{\text{ln}\left( \frac{\phi+\psi}{\phi-\psi} \right)x}{\psi}}+1}-\phi$$

Or in the cleaner form, with the substitution for $s$:

$$f(x)=\frac{2\phi e^{sx}}{e^{sx}+1}-\phi,s=\frac{\text{ln}\left( \frac{\phi+\psi}{\phi-\psi} \right)}{\psi},$$

This is the exact version we use in the project and most of the operations further down.

**S2. Algebraic construction of the benefit function**

The benefit of $n$ kilocalories to be consumed after $x$ previous kilocalories is

$$\mathrm{ben}(x,n)=\mathrm{sat}(x+n)-\mathrm{sat}(x).$$

Therefore, using the saturation function formula, we get

$$\mathrm{ben}(x,n)=\frac{2\phi e^{s(x+n)}}{e^{s(x+n)}+1}-\phi-\frac{2\phi e^{sx}}{e^{sx}+1}+\phi$$

and

$$\mathrm{ben}(x,n)=\frac{2\phi e^{s(x+n)}}{e^{s(x+n)}+1}-\frac{2\phi e^{sx}}{e^{sx}+1}.$$

Factor out $2\phi$ and convert the sum in the exponent to the product of two $e$ terms to respective powers:

$$\mathrm{ben}(x,n)=2\phi\left( \frac{e^{sx}e^{sn}}{e^{sx}e^{sn}+1}-\frac{e^{sx}}{e^{sx}+1} \right)$$

Substitute $e^{sx}$ for $a$ and $e^{sn}$ for $b$ and the bracket’s insides become:

$$\frac{ab}{ab+1}-\frac{a}{a+1}$$

Put over the common denominator:

$$\frac{a^{2}b+ab-a^{2}b-a}{(ab+1)(a+1)}$$

$$\frac{ab-a}{(ab+1)(a+1)}$$

Add $+1$ and $-1$ to the numerator (it is essentially like adding $0$ to it):

$$\frac{ab+1-a-1}{(ab+1)(a+1)},$$

which allows us to separate the fraction elegantly into two:

$$\frac{ab+1}{(ab+1)(a+1)}-\frac{a+1}{(ab+1)(a+1)}$$

leaving the expression “less wordy” after simplification:

$$\frac{1}{a+1}-\frac{1}{ab+1}$$

Which, after substituting back for a and b, is:

$$\frac{1}{e^{sx}+1}-\frac{1}{e^{sx}e^{sn}+1}$$

and, returned to the original expression with brackets, gives:

$$\mathrm{ben}(x,n)=2\phi\left( \frac{1}{e^{sx}+1}-\frac{1}{e^{sx}e^{sn}+1} \right),$$

and therefore (still with the $s$ substitution):

$$\mathrm{ben}(x,n)=\frac{2\phi}{e^{sx}+1}-\frac{2\phi}{e^{sx}e^{sn}+1},s=\frac{\text{ln}\left( \frac{\phi+\psi}{\phi-\psi} \right)}{\psi}.$$

This formula constitutes a general benefit function, which can be expressed using only the original parameters as:

$$\mathrm{ben}(x,n)=\frac{2\phi}{\left( \frac{\phi+\psi}{\phi-\psi} \right)^{\frac{x}{\psi}}+1}-\frac{2\phi}{\left( \frac{\phi+\psi}{\phi-\psi} \right)^{\frac{x+n}{\psi}}+1},$$

Since we can write

$$e^{\frac{\text{ln}\left( \frac{\phi+\psi}{\phi-\psi} \right)x}{\psi}}$$

also as

$$\left( e^{\text{ln}\left( \frac{\phi+\psi}{\phi-\psi} \right)} \right)^{\frac{x}{\psi}},$$

which collapses into

$$\left( \frac{\phi+\psi}{\phi-\psi} \right)^{\frac{x}{\psi}}.$$

**S3. Comparison of saturation functions with different ψ parameter**

If we insist that only the first Calorie is as useful as it can be, the saturation function does not change that much (Figure S1).

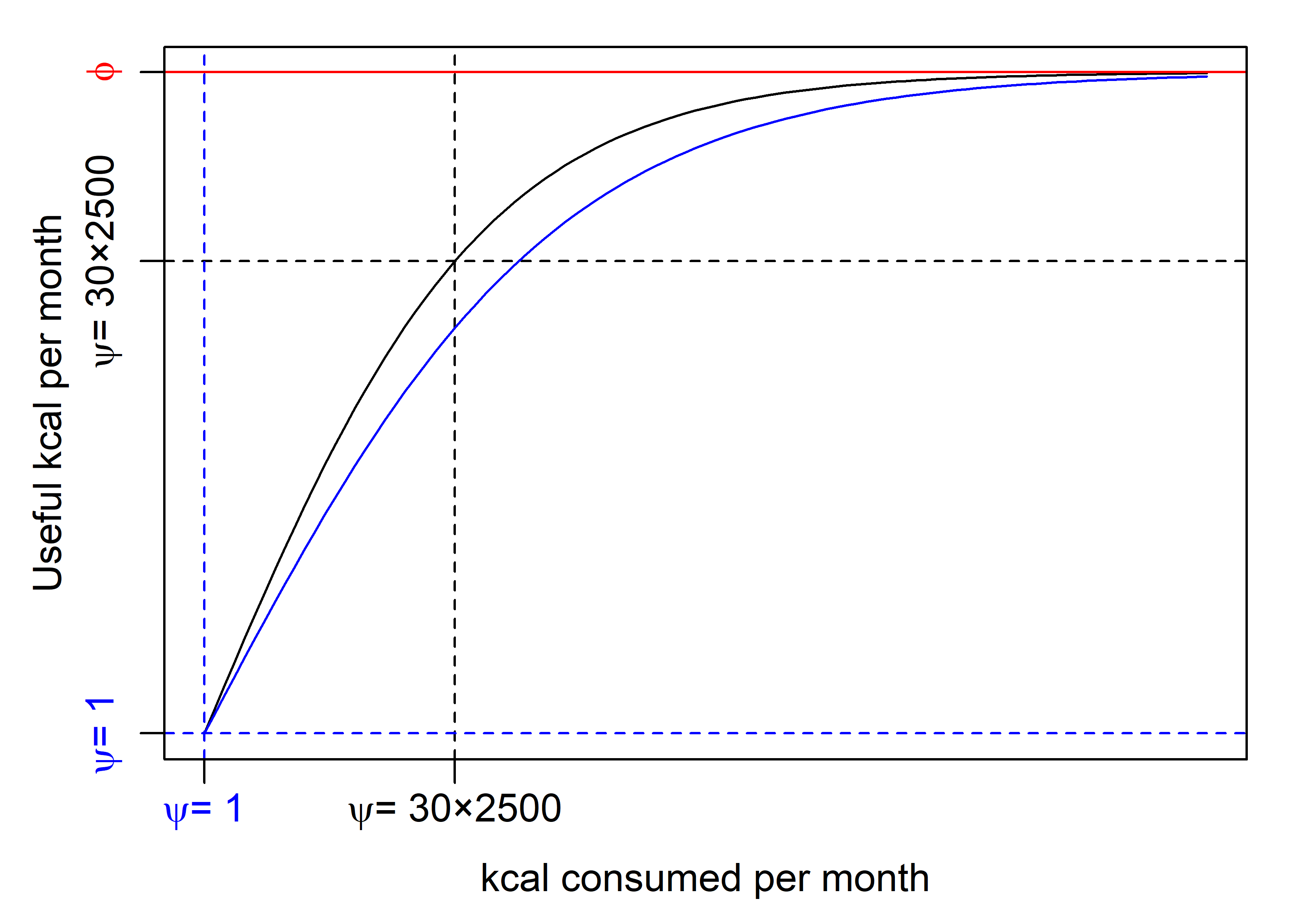

*Figure S1. Comparison of two saturation functions scaled by the decisive parameter* $\phi=3500$ *and two different parameters* $\psi=75000$ *(black curve) and* $\psi=1$ *(blue curve).*

**S4. The cost/benefit analysis - the search for a threshold**

The total expected balance of a potential new food source is a difference between expected costs and benefits:

$$\mathrm{balance}(x,n,\phi,\psi,c)=\frac{2\phi}{\left( \frac{\phi+\psi}{\phi-\psi} \right)^{\frac{x}{\psi}}+1}-\frac{2\phi}{\left( \frac{\phi+\psi}{\phi-\psi} \right)^{\frac{x+n}{\psi}}+1}-c.$$

Solving for $\mathrm{balance}(x,n,\phi,\psi,c)=0$ we get:

$$0=\frac{2\phi}{e^{st}+1}-\frac{2\phi}{e^{st}e^{sn}+1}-c,s=\frac{\text{ln}\left( \frac{\phi+\psi}{\phi-\psi} \right)}{\psi},$$

where $t$ stands for threshold, at which scrounging for the given food source is no longer profitable, $c$ is the total sum of constant costs (and possibly benefits independent of the saturation function).

At $t$, costs are balanced with benefits:

$$\frac{2\phi}{e^{st}+1}-\frac{2\phi}{e^{st}e^{sn}+1}=c$$

Substituting $a$ for $e^{st}$ we get:

$$\frac{2\phi}{a+1}-\frac{2\phi}{e^{sn}a+1}=c.$$

Add the negative bit on a left side to both sides of the equation:

$$\frac{2\phi}{a+1}=c+\frac{2\phi}{e^{sn}a+1}$$

Put over the common denominator:

$$\frac{2\phi}{a+1}=\frac{e^{sn}ca+2\phi+c}{e^{sn}a+1}$$

Multiply by $(a+1)(e^{sn}a+1)$:

$$2\phi e^{sn}a+2\phi=e^{sn}ca^{2}+2\phi a+ca+e^{sn}ca+2\phi+c$$

Subtract $2\phi e^{sn}a+2\phi$ from both sides:

$$e^{sn}ca^{2}+2\phi a+ca+e^{sn}ca-2\phi e^{sn}a+c=0$$

Factor out $a$, where convenient:

$$e^{sn}ca^{2}+(2\phi+c+e^{sn}c-2\phi e^{sn})a+c=0$$

Factor out $e^{sn}$ within the brackets:

$$e^{sn}ca^{2}+((c-2\phi)e^{sn}+2\phi+c)a+c=0,$$

which leads to a quadratic equation.

Filling the pattern:

$$a=\frac{-b\pm\sqrt{b^{2}-4\alpha c}}{2\alpha}$$

with the terms of the relationship above, we arrive at:

$$a=\frac{-((c-2\phi)e^{sn}+2\phi+c)\pm\sqrt{((c-2\phi)e^{sn}+2\phi+c)^{2}-4e^{sn}c^{2}}}{2ce^{sn}}.$$

This expression can be simplified by multiplying all terms inside the parentheses by −1, which removes one set of brackets under the square root:

$$a=\frac{(2\phi-c)e^{sn}-2\phi-c\pm\sqrt{((c-2\phi)e^{sn}+2\phi+c)^{2}-4e^{sn}c^{2}}}{2ce^{sn}},s=\frac{\text{ln}\left( \frac{\phi+\psi}{\phi-\psi} \right)}{\psi}$$

We should not forget about the substitution for $a=e^{st}$.

Therefore:

$$st=\text{ln}(a)$$

and:

$$t=\frac{\text{ln}(a)}{s}$$

The boundary of $t$ Calories, can be written as:

$$t=\frac{\text{ln}\left( \frac{(2\phi-c)e^{sn}-2\phi-c\pm\sqrt{((c-2\phi)e^{sn}+2\phi+c)^{2}-4e^{sn}c^{2}}}{2ce^{sn}} \right)}{\frac{\text{ln}\left( \frac{\phi+\psi}{\phi-\psi} \right)}{\psi}}$$

Which is:

$$t=\frac{\text{ln}\left( \frac{(2\phi-c)e^{sn}-2\phi-c\pm\sqrt{((c-2\phi)e^{sn}+2\phi+c)^{2}-4e^{sn}c^{2}}}{2ce^{sn}} \right)\psi}{\text{ln}\left( \frac{\phi+\psi}{\phi-\psi} \right)}$$

So substituting back for $s$ everywhere, the result would be:

$$t=\frac{\text{ln}\left( \frac{\left( 2\phi-c \right)\left( \frac{\phi+\psi}{\phi-\psi} \right)^{\frac{n}{\psi}}-2\phi-c\pm\sqrt{\left( (c-2\phi)\left( \frac{\phi+\psi}{\phi-\psi} \right)^{\frac{n}{\psi}}+2\phi+c \right)^{2}-4\left( \frac{\phi+\psi}{\phi-\psi} \right)^{\frac{n}{\psi}}c^{2}}}{2c\left( \frac{\phi+\psi}{\phi-\psi} \right)^{\frac{n}{\psi}}} \right)\psi}{\text{ln}\left( \frac{\phi+\psi}{\phi-\psi} \right)}$$

Technically, the expression returns two values because there are two solutions for a quadratic equation (see the $\pm$ before the discriminant). Of course, negative numbers make no sense as consumption stopping points, so only the positive real numbers should be accepted as sensible threshold solutions.

The main article’s formula does not substitute all the way back in the expression above and highlights the search for a positive threshold by keeping just the $+$ instead of $\pm$ before the discriminant; it is expressed as:

$$t=\frac{\text{ln}(a)}{s},a=\frac{(2\phi-c)e^{sn}-2\phi-c+\sqrt{((c-2\phi)e^{sn}+2\phi+c)^{2}-4e^{sn}c^{2}}}{2ce^{sn}},$$

$$s=\frac{\text{ln}\left( \frac{\phi+\psi}{\phi-\psi} \right)}{\psi},c=?.$$

The threshold $t$ above can be read as an arbitrary threshold from a continuum of differences between costs and benefits $\delta$; $t_{0}$ for $\delta=0$. Deriving the threshold for any $\delta$ over a continuum of $\rho$ values would allow analytical derivation of the isobalance contours shown in Figure 4, rather than approximating them on a discrete parameter grid.

Any contour line connecting points with equal expected payoff $\delta$ can be found using the expression:

$$t_{\delta}=\frac{\text{ln}(a)}{s},a=\frac{(2\phi-c)e^{sn}-2\phi-c\pm\sqrt{((c-2\phi)e^{sn}+2\phi+c)^{2}-4e^{sn}c^{2}}}{2ce^{sn}},$$

$$s=\frac{\text{ln}\left( \frac{\phi+\psi}{\phi-\psi} \right)}{\psi},c=\mathbb{E}[c_{D}]+\mathbb{E}[c_{A}]+e^{\left( \mu+\frac{\rho\sigma^{2}}{2} \right)}+\delta.$$

**S5. Alternative parametrisations of the log-normal distribution**

It takes two parameters to define a log-normal distribution, but they do not necessarily need to be those two in the exponentiated normal distribution $\mu$ and $\sigma$, nor the two that represent the arithmetic mean and standard deviation of the log-normally distributed variable $\mu_{X}$ and $\sigma_{X}$.

The expected value $\mathbb{E}[x]$ might be referred to as:

$$\mu_{X}=\frac{1}{n}\sum_{i=1}^{n} x_{i}$$

and the expected variance:

$$\sigma_{X}^{2}=\frac{1}{n}\sum_{i=1}^{n} (\mu_{X}-x_{i})^{2}$$

The original parameters of the distribution itself – i.e. mean and standard deviation of the normal distribution used in the exponent keep labels $\mu$ and $\sigma$.

It is mentioned in the main article, how to get $\mu_{X}$ out of $\mu$ and $\sigma$ in the expression for expected costs $\mathbb{E}[c]$.

$$\mu_{X}=e^{(\mu+\frac{\sigma^{2}}{2})}$$

Expected variance can be calculated using the following formula:

$$\sigma_{X}^{2}=(e^{\sigma^{2}}-1)e^{2\mu+\sigma^{2}}$$

Inversely, we can determine $\mu$ and $\sigma$ using:

$$\mu=\text{ln}\left( \frac{\mu_{X}^{2}}{\sqrt{\mu_{X}^{2}+\sigma_{X}^{2}}} \right)$$

and:

$$\sigma^{2}=\text{ln}\left( 1+\frac{\sigma_{X}^{2}}{\mu_{X}^{2}} \right).$$

Using the formula for $\mu_{X}$, we can easily calculate parameter $\sigma$ from mean $\mu_{X}$ and the logarithm of median $\mu$:

$$\sigma^{2}=2(\text{ln}(\mu_{X})-\mu)$$

or from the logarithm of median $\mu$ and expected variance $\sigma_{X}^{2}$:

$$\sigma^{2}=\text{ln}\left( \frac{e^{2\mu}+\sqrt{e^{4\mu}+4e^{2\mu}\sigma_{X}^{2}}}{2e^{2\mu}} \right)$$

These equations are sufficient to convert any parametrisation of log-normal distribution into any other. We can now calculate $\sigma$ from any two parameters and calculate each parameter in at least one way. In total there are, however, 12 equations that allow computing any parameter from any other two. We do not list them here for brevity, but these equations are employed in function $\text{fillLognorm}\text{()}$ that asks for any two parameters (from $\mu$, $\mu_{X}$, $\sigma$, $\sigma_{X}$) and delivers all four and the median (if the median $M=e^{\mu}$ is not inputted instead of median logarithm $\mu$), and the mode $m=e^{\left( \mu-\sigma^{2} \right)}$ on top.

The main reason for introducing this function is that we can frequently infer two parameters that cannot be readily inputted into the log-normal probability density function:

$$\text{PDF}_{\text{log-normal}}(x,\mu,\sigma)=\frac{1}{x\sigma\sqrt{2\pi}}e^{-\frac{(\text{ln}(x-\mu))^{2}}{2\sigma^{2}}}$$

whose parametrisation with logarithm median $\mu$ and logarithm standard deviation $\sigma$ does not suit our intuition. The function $\text{fillLognorm}\text{(}\text{)}$ has the capacity to convert ad-hoc intuitions into parameters of the log-normal distribution.

In the case of low acquisition costs (reported in the article), which correspond roughly to half an hour of cooking, we can infer the median, 80 kcal, and easily imagine the overall standard deviation. We set it to 40 (such that the hypothetical lower end of the scale is two standard deviations from the median, see Figure S2).

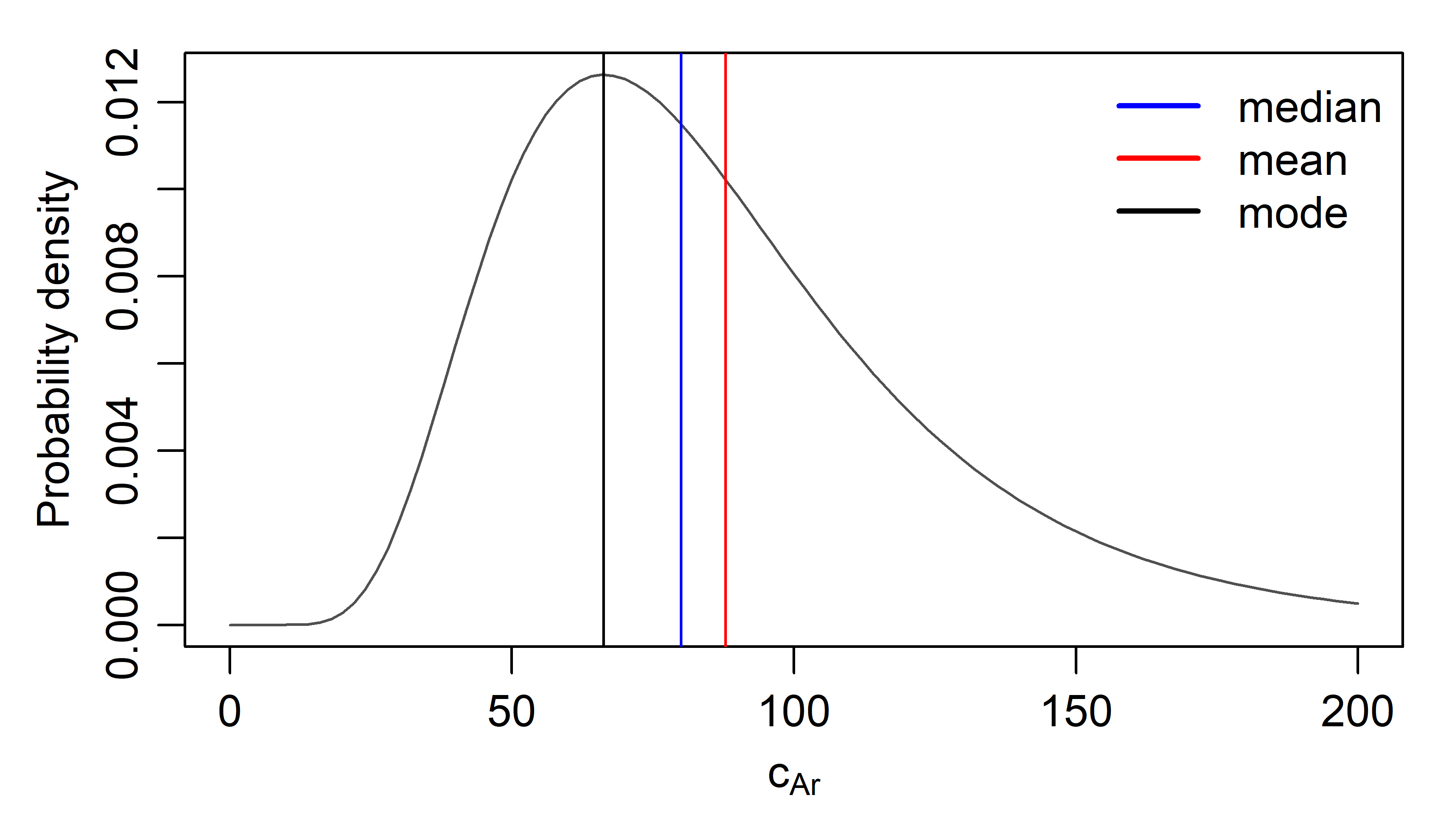

*Figure S2. Distribution of the preparation costs per portion characterised by* $M=80$ *(blue line) and* $\sigma_{X}=40$*, the* $\text{fillLognorm}\text{()}$ *function that we offer in script (https://osf.io/cmnsf) returns parameters* $\mu_{X}=87.89$ *(red line),* $\mu=4.38$*,* $\sigma=0.43$*, and* $m=66.27$ *(black line) if inputted with these two. The original parameters are also parsed in the function’s output for consistency.*

**S6. Log-normal distribution as a combination of multiplicative and additive processes**

Frequently, we need to approximate a sum of multiple numbers from various log-normal distributions by a single log-normal distribution, for instance, if we require distribution of costs over multiple portions instead of costs distribution per portion.

Formulas converting between log-normal distribution parametrisations (Supplement S5) allow inferring parameters of distributions resulting from a mix of multiplicative (e.g. distribution of digestion costs per portion) and additive (e.g. summing costs over multiple portions) processes.

Another important bit is the summing of variances of uncorrelated random variables. The distribution of sums of two uncorrelated random variables $X_{1}$ and $X_{2}$, with means $\mu_{X_{1}}$, $\mu_{X_{2}}$ and standard deviations $\sigma_{X_{1}}$, $\sigma_{X_{2}}$, is expected to have mean $\mu_{X_{1}}+\mu_{X_{2}}$ and standard deviation $\sqrt{\sigma_{X_{1}}^{2}+\sigma_{X_{2}}^{2}}$ (Fisher, 1918).

If we start with a log-normal distribution with, say, $\mu=5,\sigma=1$ (Figure S3A), we calculate $\mu_{X}=e^{(5+\frac{1^{2}}{2})}=244.69$ and $\sigma_{X}=\sqrt{(e^{1^{2}}-1)e^{2\times5+1^{2}}}=320.75$.

The distribution of sums of three numbers drawn from this distribution will have $\mu_{3X}=3\times244.69=734.08$ and $\sigma_{3X}=\sqrt{3}\times320.75=555.56$. The log-normal distribution of this sum is going to be characterised by

$$\mu_{3}=\text{ln}\left( \frac{{734.08}^{2}}{\sqrt{{734.08}^{2}+{320.75}^{2}}} \right)=6.37$$

and

$$\sigma=\sqrt{\text{ln}\left( 1+\frac{{320.75}^{2}}{{734.08}^{2}} \right)}=0.67 .$$

Results of a computer simulation (draw three numbers from the unit distribution, sum them up, repeat 100 000 times to get a sample, use kernel density estimation to approximate the distribution curve) agree perfectly with the distribution characterised by these parameters (Figure S3B).

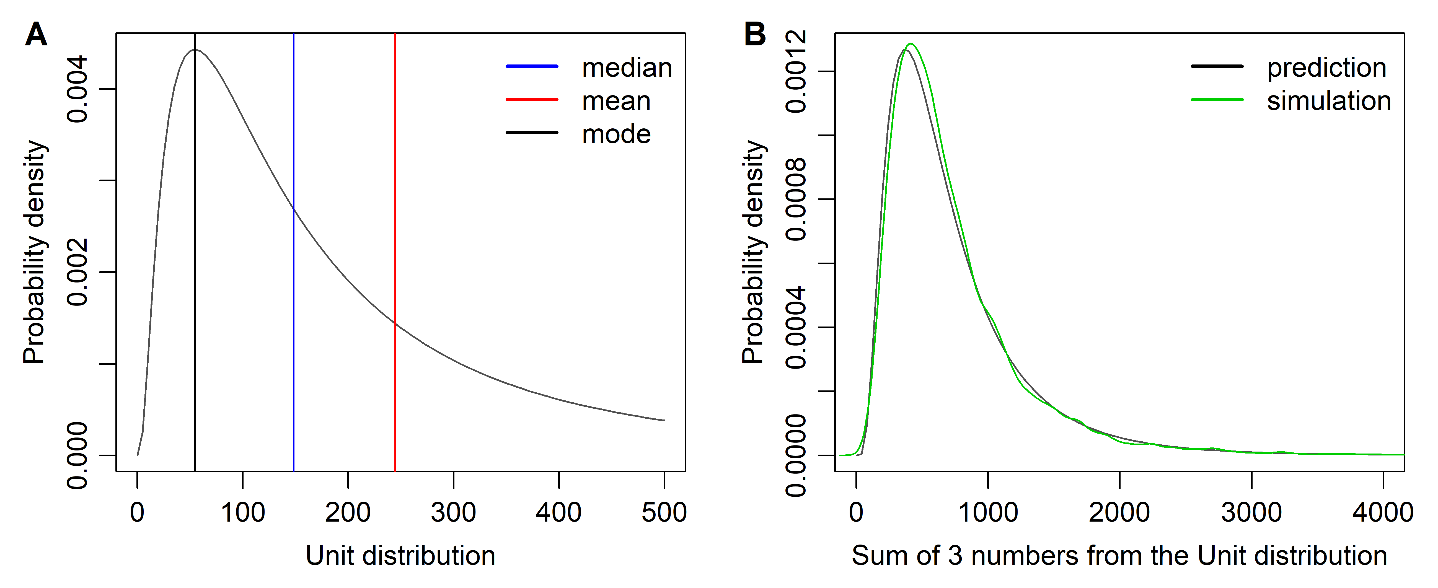

*Figure S3. The sum of numbers coming from a log-normal distribution (A) can be expressed as a log-normal distribution (B). Simulation and analytical results converge.*

We used this approach to get the whole distribution of digestion costs. The unit digestion costs distribution $c_{Dr}$ was characterised by $\mu_{Dr}=2.40$ (following from $\mu_{Dr}=\text{ln}(M_{Dr}),\mu_{Dr}=\text{ln}(11)$) and $\sigma_{Dr}=0.1$ (Figure S4A).

The expected value of this log-normal distribution is $\mathbb{E}[c_{Dr}]=11.06$ and the standard deviation $\sigma_{XDr}=1.11$. The total digestion costs in 64.75 portions of human flesh are $\mathbb{E}[c_{D}]=64.75\times11.06,\mathbb{E}[c_{D}]=715.84$. The standard deviation of this sum is $1.11\times\sqrt{64.75}=8.93)$, which, together with the arithmetic mean, leads to a log-normal distribution with parameters $\mu_{D}=6.57$ and $\sigma_{D}=0.012$. These two parameters characterise the whole distribution of digestion costs per 64.75 portions from the definition of log-normal distribution (Figure S4B). However, the mean value equal to the expected costs can be inferred without information about standard deviation and is sufficient to calculate the expected balance.

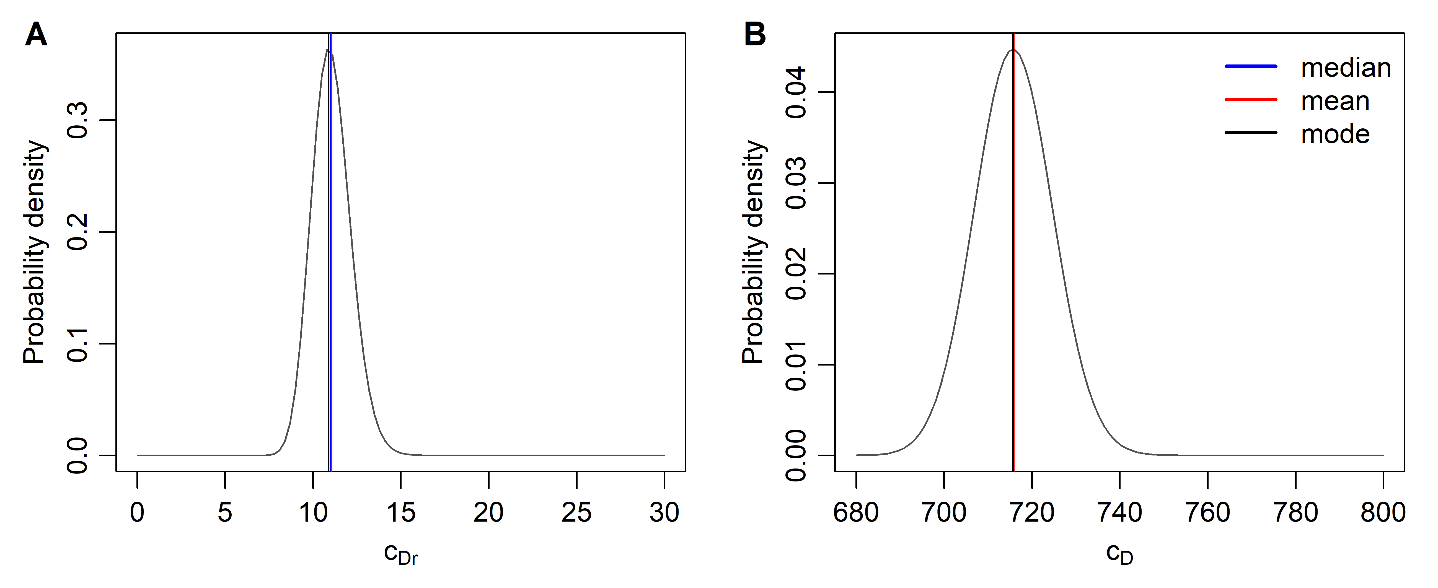

*Figure S4. Digestion costs per portion (A) and per 64.75 portions (B)*

The median food preparation cost is 80 kcal with $\sigma_{X}=40$ kcal (Figure S5A). For the high acquisition costs scenario, we found a distribution corresponding to cooking 64.75 dishes (Figure S5B) plus costs connected with the hunt (Figure S5C). Log-normal distribution with $\mu_{XA}=7\times3500+64.75\times87.89$ and $\sigma_{XA}=\sqrt{7^{2}\times{3500}^{2}+64.75\times{40}^{2}}$ is characterised by $\mu_{A}=10.06$, $\sigma_{A}=0.71$ (Figure S5D). The expected value $\mathbb{E}[c_{A}]=30191.36$, which is almost as much as the Caloric value of the human body (Figure S5D).

In the low acquisition costs, log-normal distribution with $\mu_{XA}=64.75\times87.89$ and $\sigma_{XA}=\sqrt{64.75}\times40$ is required. It is characterised by $\mu_{A}=8.64,\sigma_{A}=0.057$, the expected value $\mathbb{E}[c_{A}]=5691.36$ (Figure S5B).

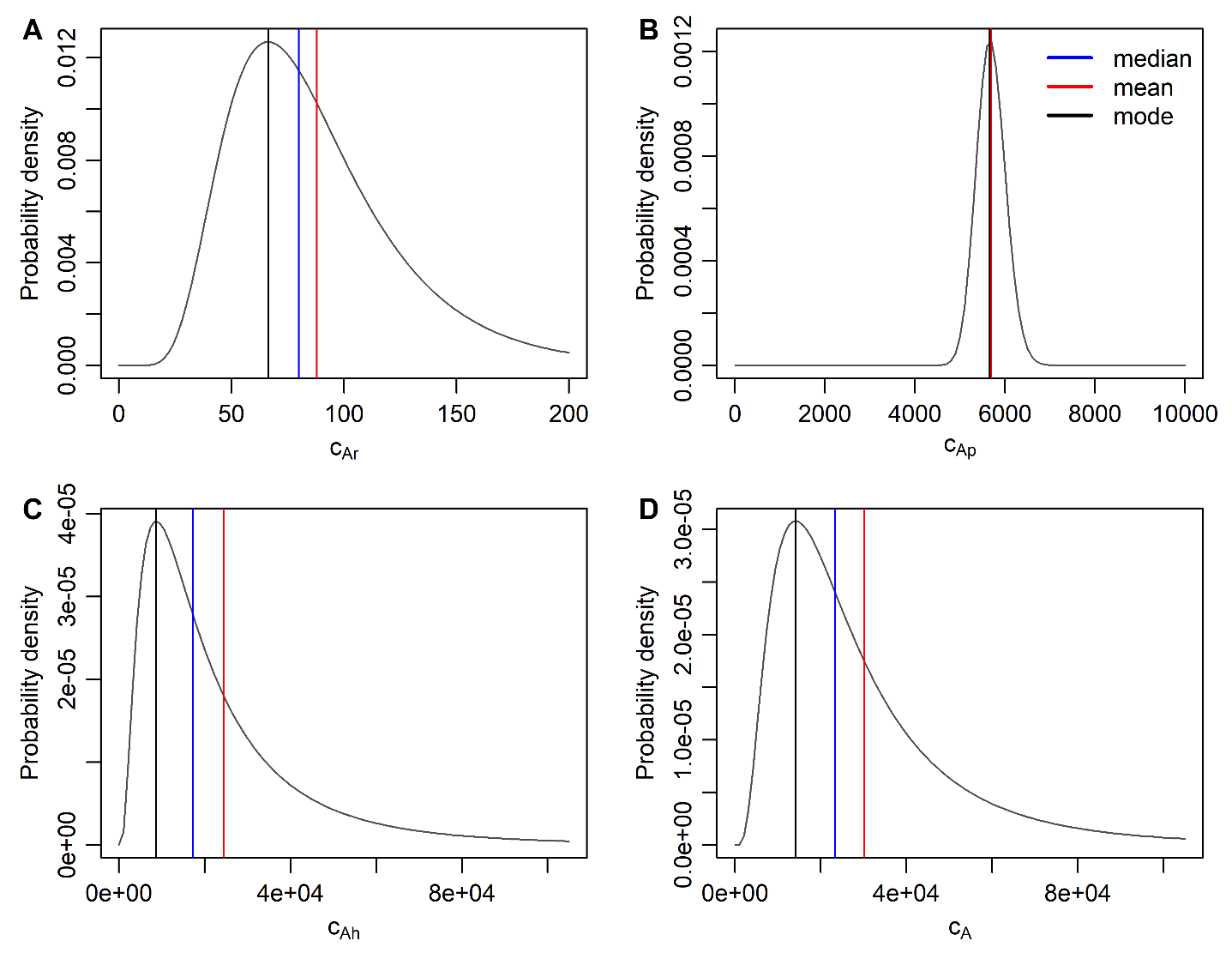

*Figure S5. Acquisition costs: Preparation per portion (A), per 64.75 portions (B), hunt for humans (C), total acquisition costs in the costly scenario including the hunt (D)*

**S7. Etudes for the mathematical model of food source utilisation**

We use a bowl of spaghetti as a benchmark food source. We will consider a unit of this source: a single bowl, allegedly 221 kcal worth in a monthly frame (Lehman, 2021).

Acquisition and preparation costs per portion correspond to the unit digestion and cooking costs in the main article. Infection costs parameters were set to $\mu_{I}=0,\sigma_{I}=0$, because spaghetti is very safe to eat.

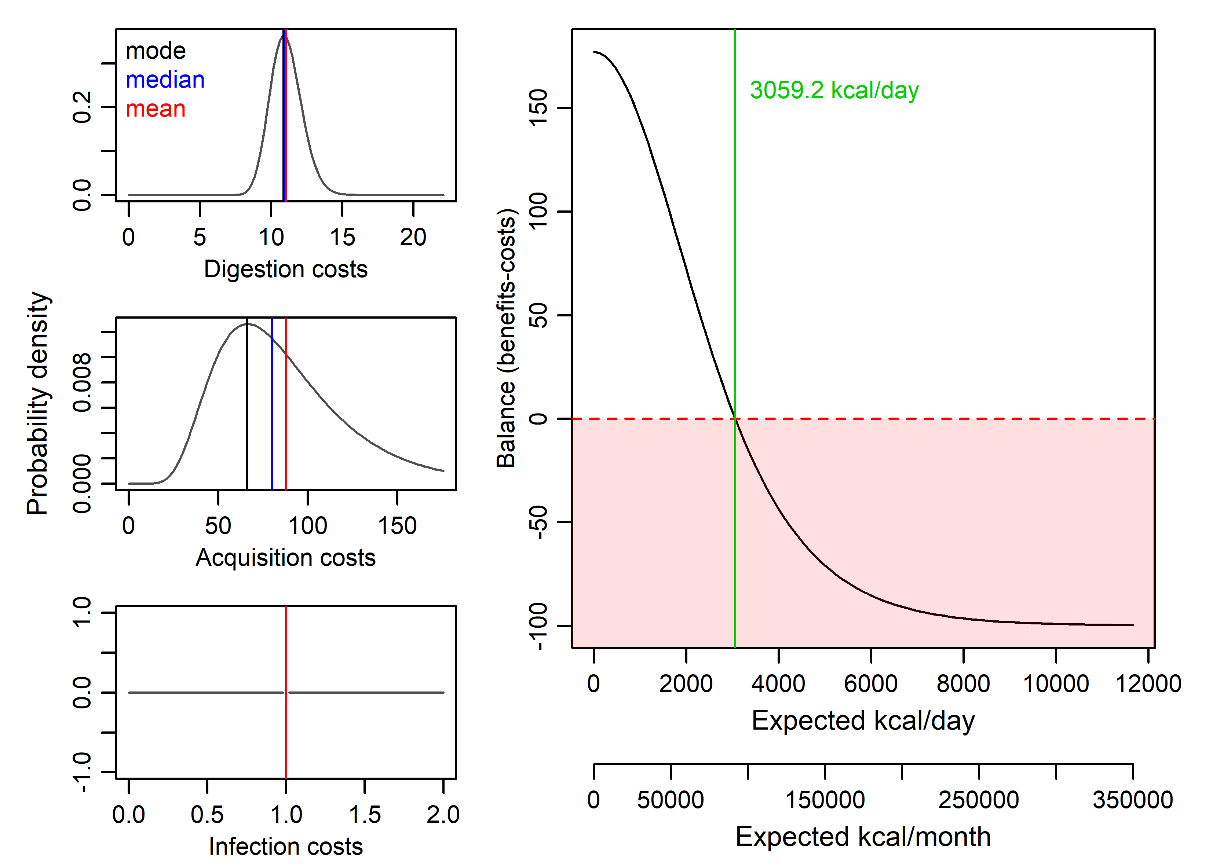

*Figure S6. Cost-benefit analysis of spaghetti consumption,* $\phi=30\times3500$*,* $\psi=30\times2500$*,* $n=221$*,* $\mathbb{E}[c_{D}]=11.06$*,* $\mathbb{E}[c_{A}]=87.89$*,*$\mathbb{E}[c_{I}]=1$*.*

The model assumes that when your daily Calorical intake is 3059.2 or more, you resign on cooking a bowl of spaghetti (Figure S6).

Grass provides a useful point of contrast. Despite its widespread availability, grass is not consumed as a food source. From the perspective of acquisition costs, grass is readily accessible and can be collected with minimal energetic expenditure. Accordingly, acquisition costs are limited primarily to basic movement. We therefore set the median acquisition costs of grass to 10 kcal, with a standard deviation of 8 kcal.

The Caloric value of a grass bowl is much lower than the spaghetti bowl, largely due to its high cellulose content. For example, one cup of mixed salad greens contains approximately 9 kcal (Fatsecret, 2021), and a typical bowl corresponds to roughly two cups (18 kcal). For the purpose of direct comparison with spaghetti, we apply the same expectations for digestion and infection costs, although processing a bowl of grass may in practice require greater digestive effort than a spaghetti dish.

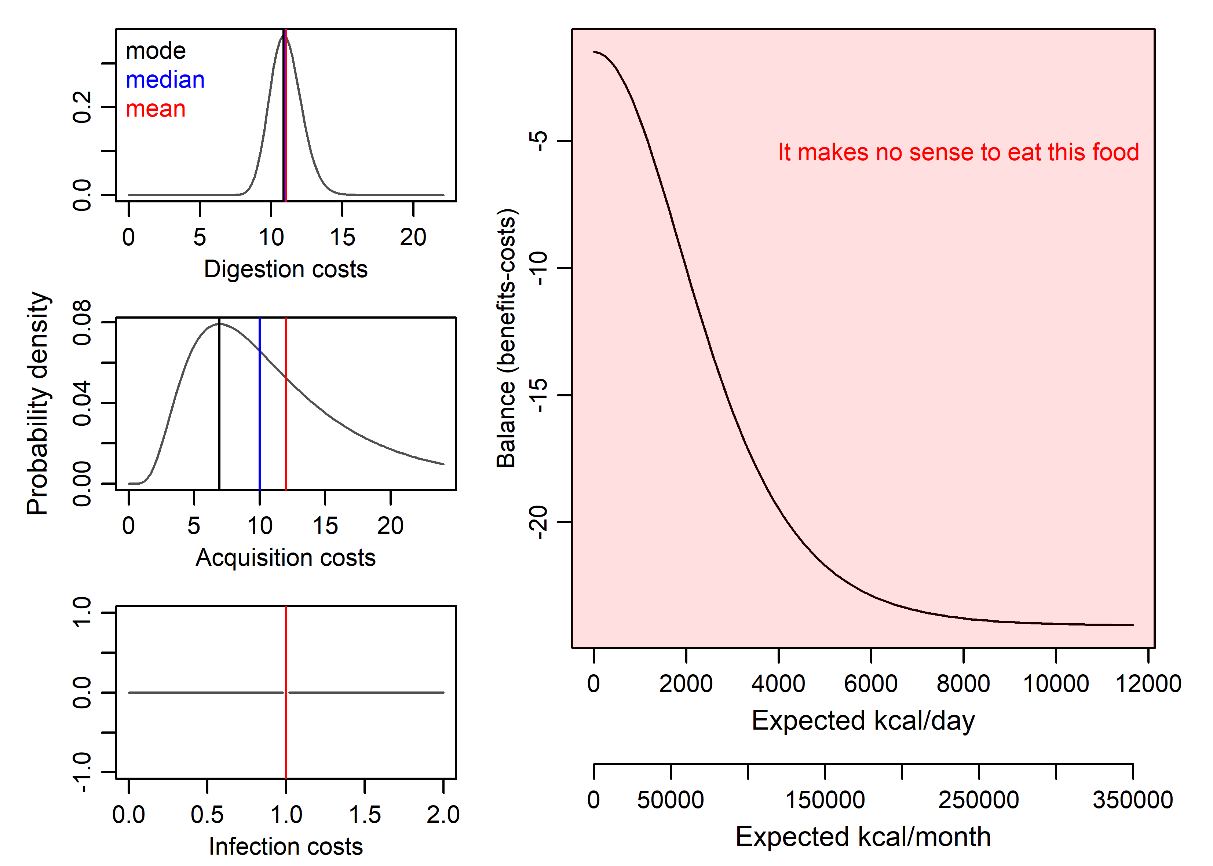

*Figure S7. Cost-benefit analysis of grass consumption,* $\phi=30\times3500$*,* $\psi=30\times2500$*,* $n=18$*,* $\mathbb{E}[c_{D}]=11.06$*,* $\mathbb{E}[c_{A}]=12.01$*,* $\mathbb{E}[c_{I}]=1$*.*

Even when the daily energetic balance is zero, grass is not incorporated into the diet because the energetic benefits are outweighed by the costs (Figure S7). This is not due to high acquisition costs, but rather to limited digestibility, which results in low effective caloric returns.

Other plant-based resources provide a contrasting case. Unlike grass, nettles can be processed into edible dishes, for example by boiling them into a soup (see, e.g. Tanner, 2021). This additional processing increases acquisition costs, while digestion and infection costs are held constant for comparability. Under these assumptions, a serving of nettles can yield a substantial fraction of the caloric content of a spaghetti dish. Although the resulting energetic return remains modest, there are conditions under which nettles provide a higher expected energetic balance than consuming no food.

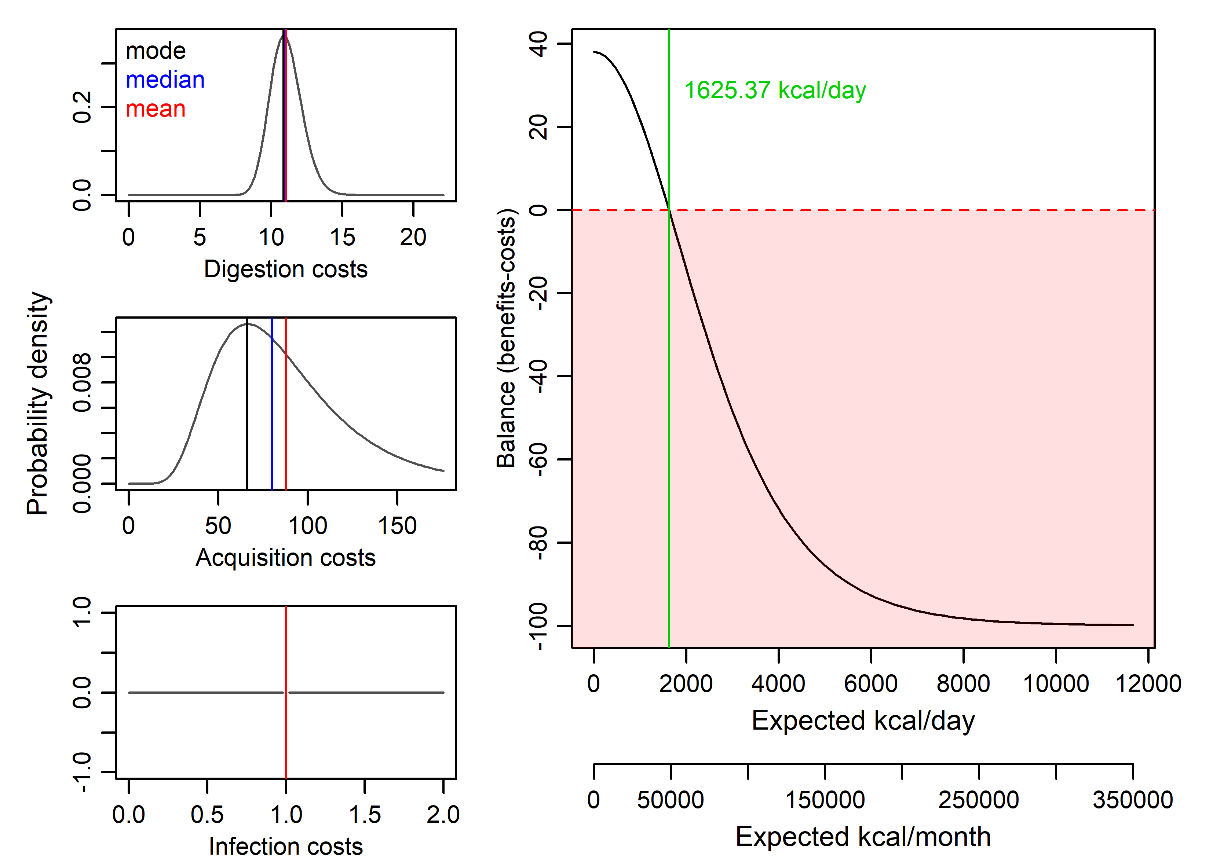

*Figure S8. Cost-benefit analysis of nettle-soup consumption,* $\phi=30\times3500$*,* $\psi=30\times2500$*,* $n=110$*,* $\mathbb{E}[c_{D}]=11.06$*,* $\mathbb{E}[c_{A}]=87.89$*,*$\mathbb{E}[c_{I}]=1$*.*

Accordingly, nettles are not routinely incorporated into the diet under conditions of adequate food availability, as alternative food sources with higher energetic returns are accessible. However, when expected daily caloric intake falls below 1625.37 kcal, the model predicts that nettles yield a higher expected energetic balance than consuming no additional food, making their consumption energetically advantageous under conditions of scarcity (Figure S8).

Thus far, the analysis has focused on plant-based food sources. As a baseline comparison involving animal-based food, we consider chicken, one of the most commonly consumed vertebrate food sources in contemporary human diets. The caloric value of a chicken is estimated at 1037 kcal.

Digestion and acquisition costs are assumed to correspond to the digestion and preparation of two portions, reflecting relatively low acquisition costs associated with obtaining a farmed chicken. Infection costs are assumed to be low if the meat is consumed soon after slaughter and cooked properly. Accordingly, we set infection costs to a baseline log-normal distribution with parameters $\mu_{I}=1$ and $\sigma_{I}=1$ (Figure S9).

We anchor other Infection costs approximations in this baseline. We expect mammals to host roughly twice as many pathogens viable in the human organism than birds, so we employ $\mu_{I}=2,\sigma_{I}=\sqrt{2}$ for mammals. And humans might be, when it comes to Infection costs, twice as dangerous food source as an average mammal, so we approximate first-order cannibalism with $\mu_{I}=4,\sigma_{I}=2$.

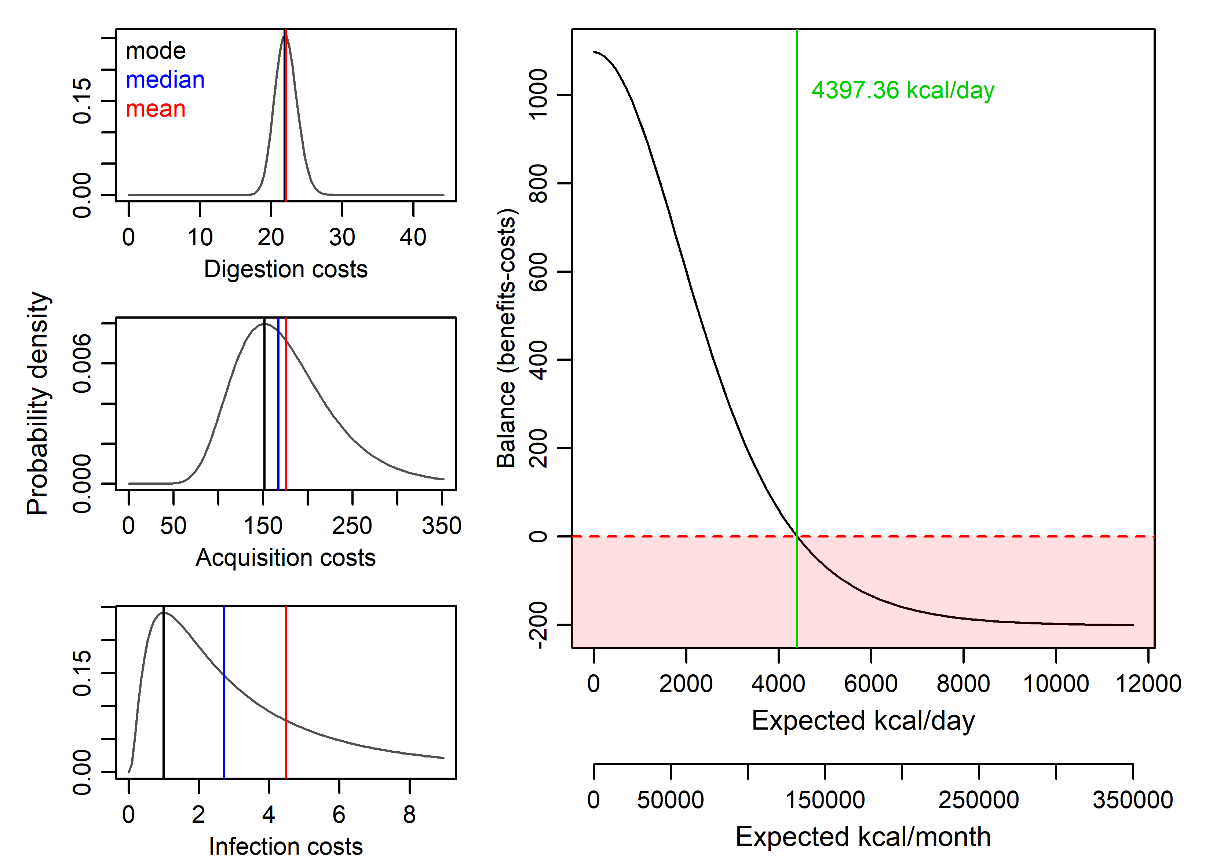

*Figure S9. Cost-benefit analysis of chicken consumption,* $\phi=30\times3500$*,* $\psi=30\times2500$*,* $n=1037$*,* $\mathbb{E}[c_{D}]=2\times11.06$*,* $\mathbb{E}[c_{A}]=2\times87.89$*,*$\mathbb{E}[c_{I}]=4.48$*.*

As a further comparison, we consider mammoth as a high-caloric animal-based food source historically available to human populations. Based on estimates of caloric content in animals potentially hunted by early humans (Cole, 2017), the total caloric value of a mammoth is approximately 3.6 million kilocalories. Given constraints on preservation and consumption, it is assumed that a mammoth could be consumed over a period of approximately one month before spoilage, and we therefore retain a 30-day reference frame.

Digestion costs are calculated based on the number of portions that can realistically be processed within this time window, rather than on the total number of potential portions contained in the mammoth. Using a daily maximum effective caloric intake of $\phi=3500$ kcal and an approximation of 500 kcal per portion, the maximum digestible intake corresponds to seven portions per day, or 210 portions per month. Accordingly, digestion costs are scaled to this number of portions.

To maintain consistency with these constraints, the energetic benefits are capped at 105,000 kcal, corresponding to the caloric content of 30 days of consumption at seven portions per day. Using the full caloric value of the mammoth would yield qualitatively similar results and does not materially affect the conclusions.

Acquisition costs are approximated using an upper-bound energetic reference corresponding to mortality. Given that a mammoth posed a lethal threat to an isolated human hunter, the expected acquisition cost is set to the energetic equivalent of the loss of a single human life. To reflect variability in hunting outcomes, the standard deviation is set equal to the mean, capturing both rare successful kills by solitary hunters and more frequent scenarios in which multiple hunters are killed during the attempt.

As mammoths were herbivores, infection costs are assumed to be relatively low. However, because mammoths are mammals, we approximate infection costs as corresponding to twice the baseline pathogen load assumed for chicken.

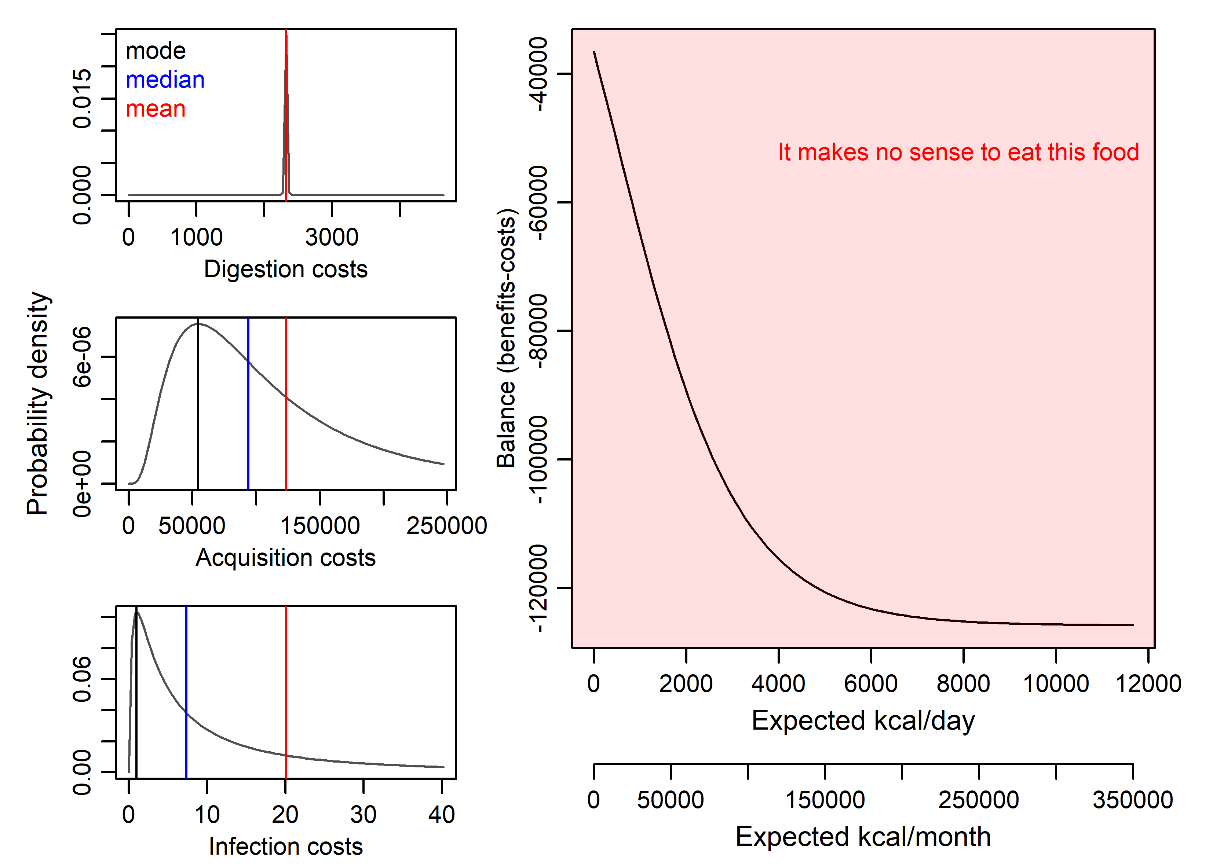

*Figure S10. Cost-benefit analysis of mammoth consumption,* $\phi=30\times3500$*,* $\psi=30\times2500$*,* $n=105000$*,* $\mathbb{E}[c_{D}]=210\times11.06$*,* $\mathbb{E}[c_{A}]=123457.9$*,* $\mathbb{E}[c_{I}]=20.09$*.*

Under these assumptions, solitary hunting of mammoths yields a strongly negative expected energetic balance, even in the absence of alternative food sources (Figure S10), due to the high probability of lethal outcomes. In such cases, mammoth hunting is not energetically viable at the individual level.

In contrast, cooperative hunting alters the expected energetic balance. When the total caloric value of approximately 3,600,000 kcal is distributed among multiple hunters (e.g., ten individuals), the energetic benefits per hunter remain substantial relative to individual consumption capacity, while acquisition costs are shared among participants (Figure S11). Infection costs, however, remain unchanged at the individual level, as consumption of even a single portion is sufficient to transmit pathogens carried by the mammoth.

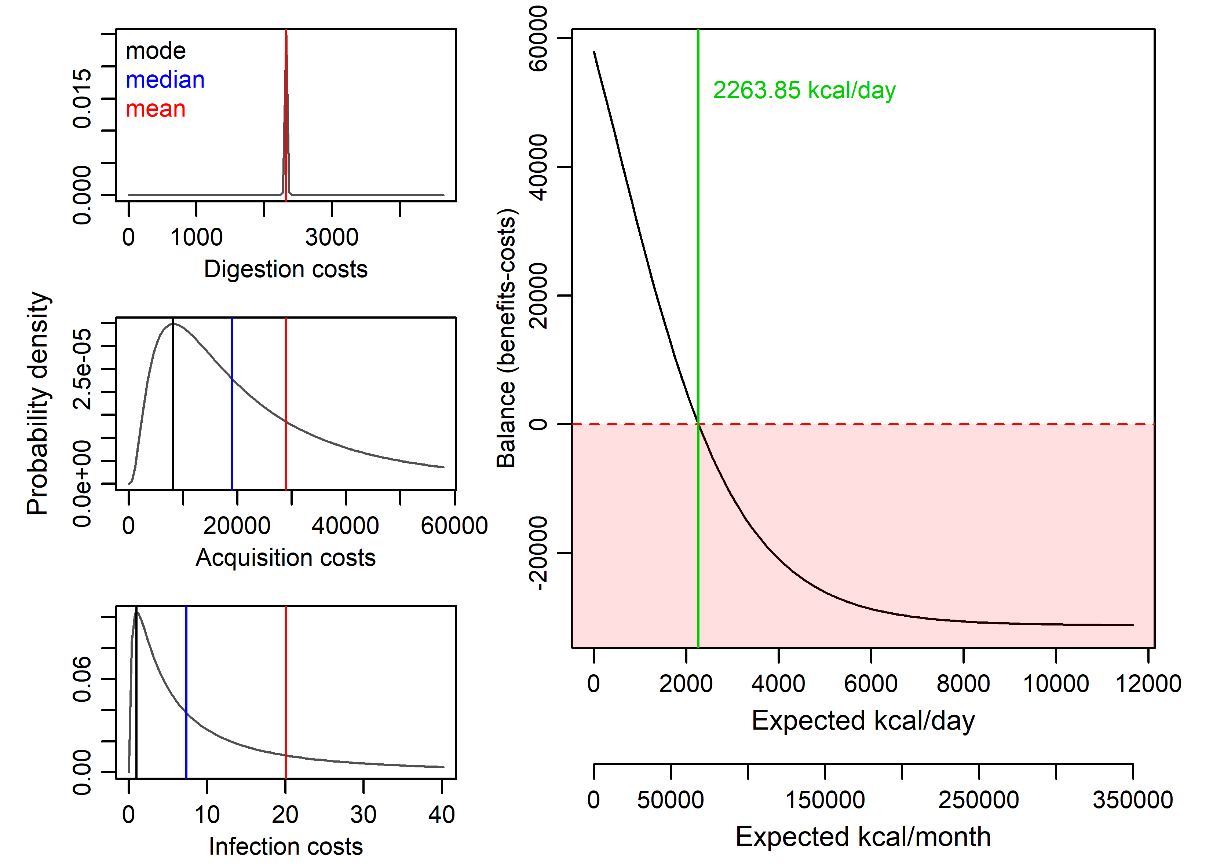

*Figure S11. Cost-benefit analysis of mammoth hunt in a group of 10 hunters,* $\phi=30\times3500$*,* $\psi=30\times2500$*,* $n=105000$*,* $\mathbb{E}[c_{D}]=210\times11.06$*,* $\mathbb{E}[c_{A}]=28957.9$*,* $\mathbb{E}[c_{I}]=20.09$*.*

Cooperation can get even more effective. In a group of 50 hunters (Figure S12), you do not reach the monthly capacity - there are going to be no leftovers. This means 72000 kcal, i.e. 144 portions of food for each hunter involved.

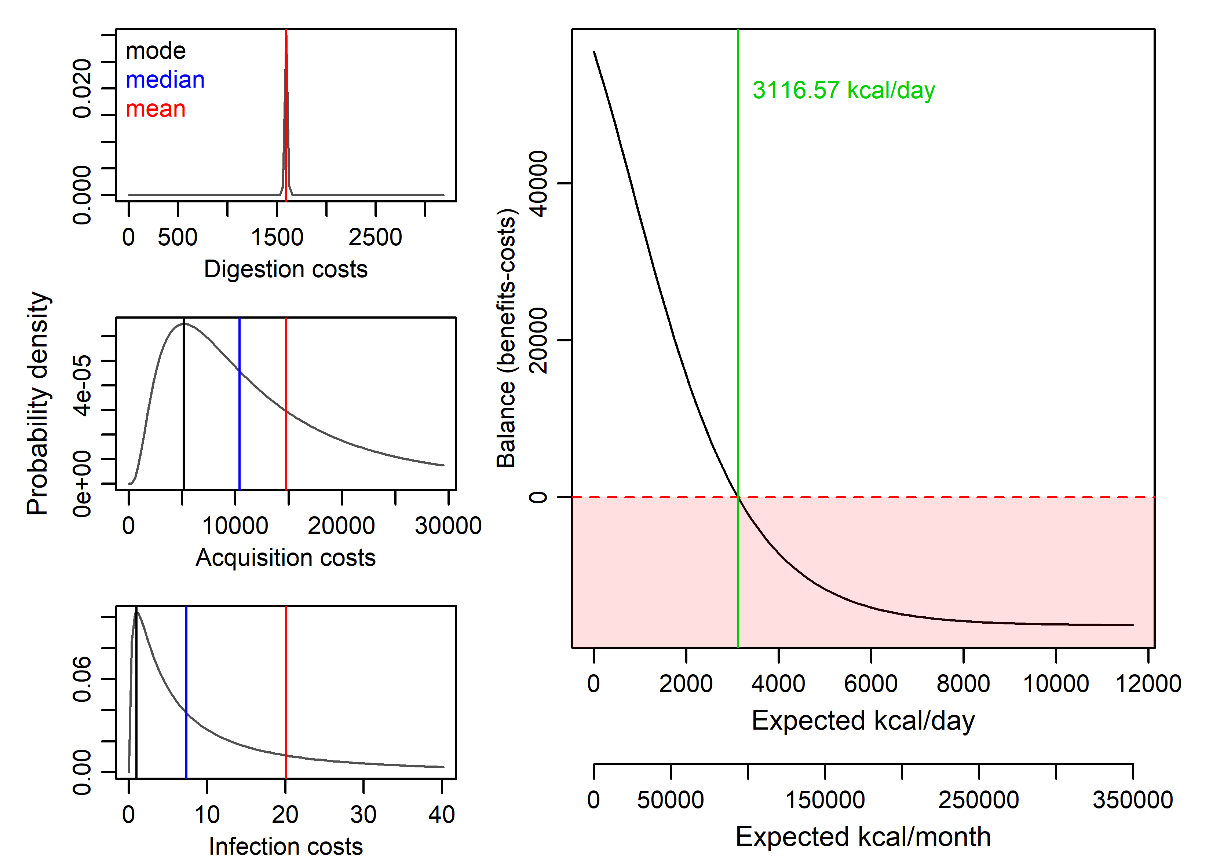

*Figure S12. Cost-benefit analysis of mammoth hunt in a group of 50 hunters,* $\phi=30\times3500$*,* $\psi=30\times2500$*,* $n=72000$*,* $\mathbb{E}[c_{D}]=144\times11.06$*,* $\mathbb{E}[c_{A}]=14756.8$*,* $\mathbb{E}[c_{I}]=20.09$*.*

This strategy is very profitable (Figure S12). Until the expected daily intake crosses 3116.57 kcal, it pays to go hunt for mammoths in groups of 50 men (each of them faces a chance of about 1.5 in 1000 that he does not survive the hunt).

Carnivorous prey provides a further point of comparison. As an illustrative case, we consider a lion. Following (Cole, 2017), we approximate a lion as energetically comparable to a 100 kg reindeer, yielding an estimated caloric value of approximately 60,000 kcal. Relative to a mammoth, a lion is assumed to pose a lower acquisition risk; accordingly, acquisition costs are set at approximately half those associated with mammoth hunting.

As a mammalian carnivore, a lion is assumed to carry a higher pathogen load than herbivorous prey. Infection costs are therefore parameterised as $\mu_{I(\text{lion})}=\mu_{I(\text{mammoth})}$ and $\sigma_{I(\text{lion})}=\sqrt{2}\text{ }\sigma_{I(\text{mammoth})}$, reflecting a trophic chain position of $\rho=2$, given that lions predominantly consume herbivores.

As in the mammoth case, the lion hunt is modelled as a cooperative activity involving ten hunters, with acquisition costs divided equally among participants.

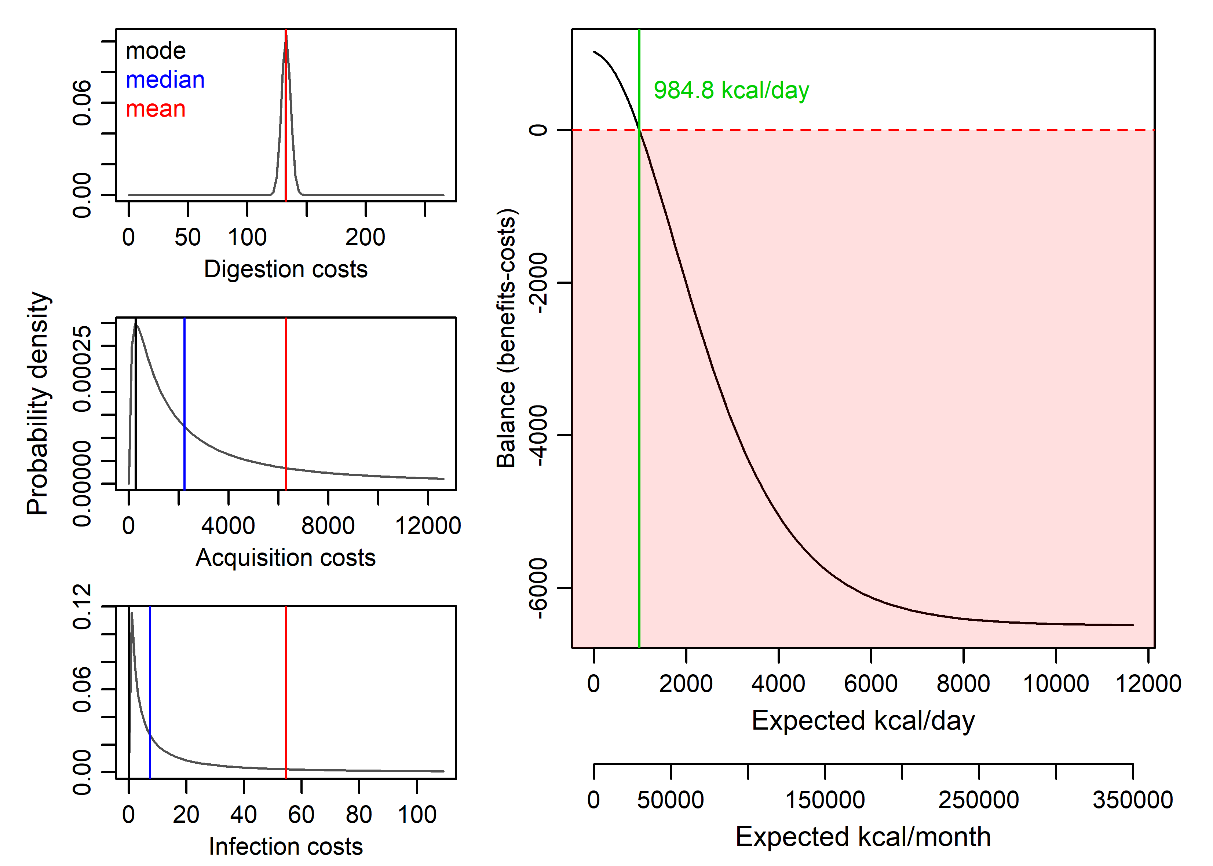

*Figure S13. Cost-benefit analysis of lion hunt in a group of 10 hunters,* $\phi=30\times3500$*,* $\psi=30\times2500$*,* $n=6000$*,* $\mathbb{E}[c_{D}]=12\times11.06$*,* $\mathbb{E}[c_{A}]=6304.7$*,* $\mathbb{E}[c_{I}]=54.59$*.*

Lion hunting is unlikely to be energetically viable under ordinary conditions and is therefore expected to occur primarily under critical circumstances. However, this assessment changes when additional benefits beyond direct energetic returns are considered. In particular, lion hunting may function as a form of costly signalling, generating social or reputational benefits that are external to subsistence. The act of killing a lion constitutes a costly signal precisely because, for an average human male, it entails substantial risk and would be energetically irrational in the absence of such external benefits.

**S8 Extreme scenarios with** $\boldsymbol{x}\boldsymbol{=}\boldsymbol{0}$

Under conditions of complete absence of alternative food sources ($x=0$), the model predicts that consumption of conspecific meat yields a positive expected energetic balance even at high orders of cannibalism. For example, when the available individual is a third-order cannibal, raw consumption results in a positive energetic balance with a probability of 91.6%, despite the associated acquisition and infection costs (Figure S14).

Even at extreme trophic chain lengths, the expected balance remains positive in probabilistic terms: consumption of a 999th-order cannibal yields a positive net energetic balance with a probability of 53.5%. These probabilities are calculated using the cumulative distribution function specified below.

The same function is used throughout the manuscript to estimate probabilities of adverse outcomes, including mortality (defined as energetic costs exceeding $30\times3500$ kcal) and severe illness (defined as energetic costs exceeding $7\times3500$ kcal).

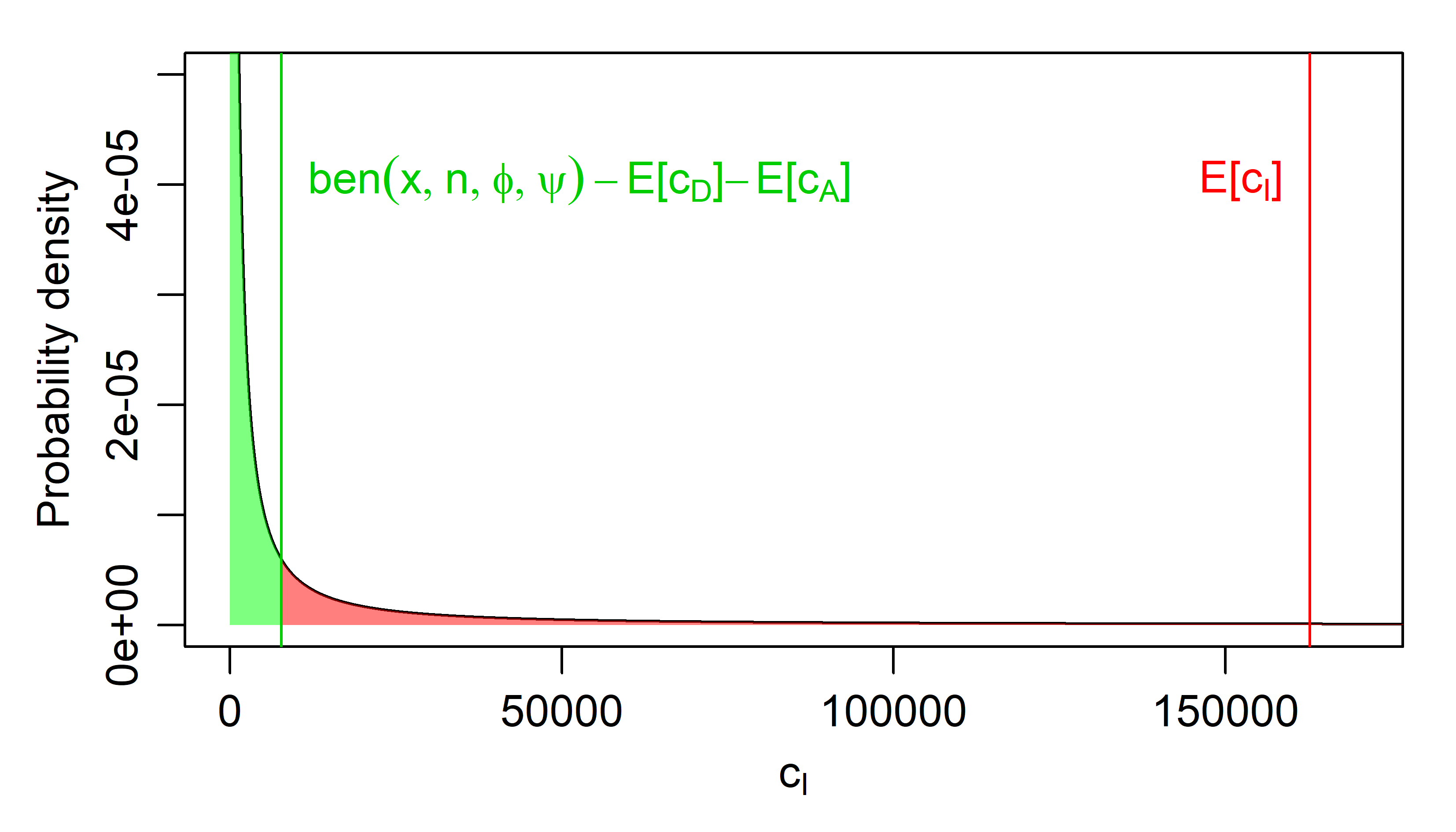

*Figure S14. Distribution of infection costs (under the assumption of no heat treatment,* $\sigma_{I}=4, \mu_{I}=2$*, and prior prey’s cannibalism* $\rho=4$*) in the context of expected benefits and other costs,* $x=0$*,* $n=32376,$ $\phi=30\times3500$*,* $\psi=30\times2500$*,* $\mathbb{E}[c_{D}]=715.84$*,* $\mathbb{E}[c_{A}]=30191.36$ *(hunt for humans)*

We consider a limiting case  corresponding to an extreme case in which a single individual encounters a recently deceased conspecific under conditions of complete isolation and absence of alternative food sources. (The “desert-island scenario”; just a lone survivor and a freshly washed up body of her colleague – a high-order cannibal perhaps.) Although the expected net energetic balance in this scenario might be strongly negative (see main text), this result is driven by the extreme right skew of the log-normal infection cost distribution at high values of $\sqrt{\rho}\sigma_{I}$. The relevant question in this context is therefore probabilistic rather than expectation-based: specifically, what is the probability that becoming a $\rho$th-order cannibal yields a positive energetic outcome in a single, isolated event.

To address this question, we reconsider the $\left( x,\rho\right)$ parameter space with explicit reference to the distribution of infection costs. Rather than analysing the expected value of the overall energetic balance, we compute the cumulative probability that consumption of a single human body results in a positive net energetic balance (Figure S14).

The cumulative distribution function of the log-normal distribution

$$\text{CDF}_{\text{log-normal}}(x,\mu,\sigma)=\frac{1}{2}+\frac{1}{2}\text{erf}\left[ \frac{\text{ln}(x)-\mu}{\sqrt{2}\sigma} \right],$$

where $\text{erf}$ represents Gauss error function

$$\text{erf}(z)=\frac{2}{\sqrt{\pi}}\int_{0}^{z} e^{-w^{2}}dw,$$

can be used to calculate the probability $q$ that the overall balance is positive, i.e. that the infection costs do not cross the difference between the expected benefits given by $\phi$, $\psi$, $n$, and $x$ and expected digestion an acquisition costs $\mathbb{E}[c_{D}]+\mathbb{E}[c_{A}]$.

The formula is

$$q=\frac{1}{2}+\frac{1}{2}\text{erf}\left[ \frac{\text{ln}(\mathrm{ben}(x,n,\phi,\psi)-\mathbb{E}[c_{D}]-\mathbb{E}[c_{A}])-\mu_{I}}{\sqrt{2\rho}\sigma_{I}} \right],$$

where $\mu_{I}$ and $\sigma_{I}$ are parameters of the first-order Infection costs distribution, $\rho$ is the cannibalism order, $x$ is the expected amount of kcal, $n$ is the Caloric value of the human body, $\phi$ is the saturation maximum, and $\psi$ is the amount of kcal as effective as measured, $\mathbb{E}[c_{D}]$ is the expected value of digestion costs, and $\mathbb{E}[c_{A}]$ is the expected acquisition costs.

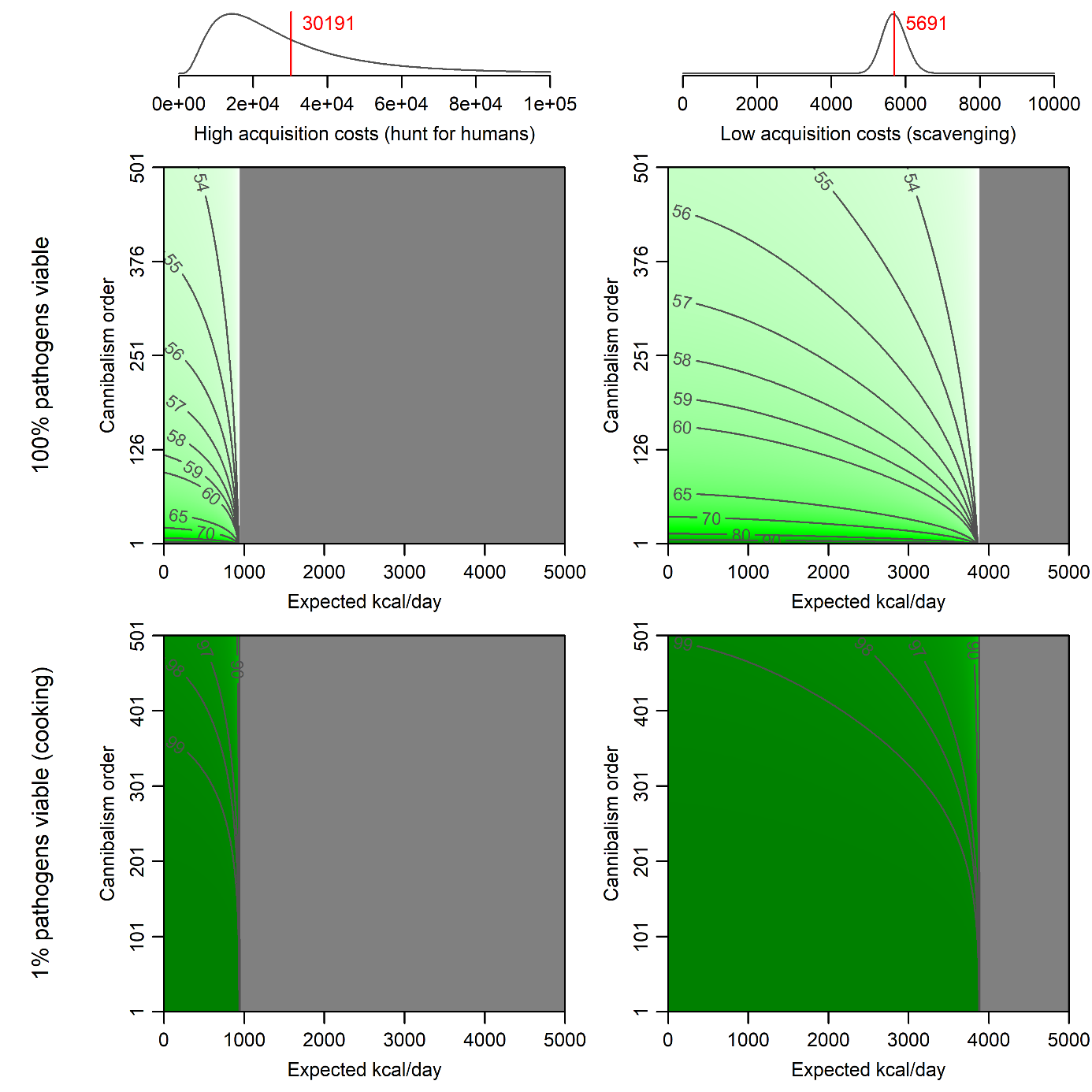

*Figure S14. Contour plot indicating the probability that a single incident of human body consumption benefits the cannibal according to food abundance and cannibalism order in two distinct levels of acquisition and infection costs. Green tones correspond to probability > 50%, grey tones to probability < 50%. Digestion costs are held constant at 11.05 kcal per 500 kcal portion of human meat. The profitability calculations are executed in a monthly frame, so the saturation function scaling parameters are* $\phi=30\times3500$*,* $n=32376$ *kcal. Infection cost of 100% pathogens are characterised by the log-normal distribution* $\mu_{I}=4,\sigma_{I}=2$*. The top and bottom row’s* $y$ *axes are identical.*

Unless the digestion and acquisition costs alone outweigh the benefits (the grey area, where the probability of profitability is zero), the consumption of the human body remains beneficial rather that detrimental. The difference between cannibalism as an isolated occasion and the long-lasting subsistence strategy is caused by the escalating discrepancy between Infection costs distribution median (stays constant) and mean (grows exponentially with the cannibalism order $\rho$.

**References**

Cole, J. (2017). Assessing the calorific significance of episodes of human cannibalism in the Palaeolithic. *Nature Publishing Group*, (April), 1–10. https://doi.org/10.1038/srep44707

Fatsecret. (2021). Calories in Salads. Retrieved March 7, 2021, from https://www.fatsecret.com/calories-nutrition/food/salad

Fisher, R. A. (1918). The Correlation between relatives on the supposition of mendelian inheritance. *Transactions of the Royal Society of Edinburgh*, (52), 399–433.

Lehman, S. (2021). Spaghetti Nutrition Facts. Retrieved March 7, 2021, from https://www.verywellfit.com/is-pasta-bad-for-your-health-2506879

Tanner, L. (2021). 10 Ingenious Foods People Ate During Famines. Retrieved March 7, 2021, from https://www.survivopedia.com/10-ingenious-foods-people-ate-during-famines/
